# Preservation of co-expression defines the primary tissue fidelity of human neural organoids

**DOI:** 10.1101/2023.03.31.535112

**Authors:** Jonathan M. Werner, Jesse Gillis

## Abstract

Human neural organoid models offer an exciting opportunity for studying often inaccessible human-specific brain development; however, it remains unclear how precisely organoids recapitulate fetal/primary tissue biology. Here, we characterize field-wide replicability and biological fidelity through a meta-analysis of single-cell RNA-sequencing data for first and second trimester human primary brain (2.95 million cells, 51 datasets) and neural organoids (1.63 million cells, 130 datasets). We quantify the degree to which primary tissue cell-type marker expression and co-expression are recapitulated in organoids across 12 different protocol types. By quantifying gene-level preservation of primary tissue co-expression, we show neural organoids lie on a spectrum ranging from virtually no signal to co-expression near indistinguishable from primary tissue data, demonstrating high fidelity is within the scope of current methods. Additionally, we show neural organoids preserve the cell-type specific co-expression of developing rather than adult cells, confirming organoids are an appropriate model for primary tissue development. Overall, quantifying the preservation of primary tissue co-expression is a powerful tool for uncovering unifying axes of variation across heterogeneous neural organoid experiments.

## Introduction

Pluripotent stem cells create self-organized multi-cellular structures, termed organoids, when cultured in a 3D *in vitro* environment^1,2^. The advantage of organoid models over 2D cell culture counterparts is their ability to generate structures that resemble endogenous tissues both in the differentiated cell-types produced and their 3D spatial organization^3,4^. The ability to model organogenesis in a controlled *in vitro* environment creates opportunities to study previously inaccessible developmental tissues from both humans and a range of model organisms^5,6,7^. As such, organoids are genetically accessible^8^ and environmentally perturbable^9^ models enabling the study of molecular, cellular, and developmental mechanisms behind tissue construction. However, the applicability of studies in organoids to *in vivo* biology hinges on how well these *in vitro* models recapitulate primary tissue developmental processes, which remains an open question.

Quantifying the degree to which organoid systems replicate primary tissue biological processes is a critical step toward understanding the strengths and limitations of these *in vitro* models^10–14^. However, studies that perform such primary tissue/organoid comparisons are inherently confounded by batch^15^ (*in vivo* vs *in vitro*), making it difficult to disentangle batch effects from underlying primary tissue and organoid biology. Meta-analytic approaches across many primary tissue and organoid datasets offer a route around these confounds, enabling the discovery of replicable primary tissue and organoid signatures independent of batch, which can then be interrogated for how well organoids recapitulate primary tissue biology^16–18^. An important biological signature for this purpose is gene co-expression^19^. Genes that are functionally related tend to be expressed together, resulting in correlated gene expression dynamics that can define functionally relevant gene modules^19^. Gene co-expression relationships represent a shared genomic space that can be aggregated across experiments (e.g.,^20^) in either *in vivo* or *in vitro* systems, thus providing a useful framework for quantifying functional similarities and differences. Excitingly, coupling meta-analytic comparisons of primary tissue and organoid co-expression with single-cell RNA-sequencing data (scRNA-seq) stands to deliver cell-type specific quantifications of organoids’ current capacity for producing functionally equivalent cell-types to primary tissues^21,22^.

Among organoid systems, human neural organoids are particularly well suited for meta-analytic evaluation due to well-described broad cell-type annotations and their known lineage relationships^23^, the wide variety of differentiation protocols in use^24^, and the increasing amount of single-cell primary brain tissue and neural organoid data publicly available. In particular, the diversity of differentiation protocols for human neural organoids poses a unique challenge for organoid quality control that can be met by meta-analytic approaches. Neural organoids can either be undirected^25^ (multiple brain region identities) or directed (specific brain region identity) with an increasing number of protocols striving to produce a wider variety of region-specific organoids^11,26–37^. Meta-analytic primary tissue/organoid comparisons across differentiation protocols stand to derive generalizable quality control metrics applicable to any differentiation protocol, fulfilling a currently unmet need for unified quality control metrics across heterogeneous neural organoids.

Prior comparisons between primary brain tissues and neural organoids demonstrated that organoids have the capacity to produce diverse cell-types that capture both regional and temporal variation similar to primary tissue data as assayed through transcriptomic^10,11,13,16,17,38^, epigenomic^39,40^, electrophysiologic^41^, and proteomic studies^42^. At the morphological level, neural organoids can produce cellular organizations structurally similar to various *in vivo* brain regions, including cortical layers^43^ and hippocampus^27^, as well as modeling known inter-regional interactions like neuromuscular junctions^34^ and interneuron migration^29^. Additionally, several prior studies have compared primary tissue/organoid co-expression and concluded that neural organoids recapitulate primary brain tissue co-expression^5,13,39^, but these assessments are highly targeted to study-specific properties, limiting potential generalization or potential assessment across the field. Typically, only a single organoid differentiation protocol is used in these assessments and it remains unclear whether organoids across different protocols will produce similar results. This lack of breadth also affects the use of primary tissue data used as a reference, with the primary tissue datasets utilized being treated as gold-standard datasets with little consideration for the extent one primary tissue reference may generalize to another. While prior meta-analytic comparisons of primary tissue/organoid co-expression have been performed^17^, these were done at the bulk level (lack cell-type resolution) and included a small number of cortical organoid protocols, limiting the biological resolution and generalizability of these findings.

In this study, we perform a meta-analytic assessment of primary brain tissue (2.95 million cells, 50 datasets, Fig. 1A) and neural organoid (1.63 million cells, 130 datasets, 12 protocols, Fig. 1B) scRNA-seq datasets, constructing robust primary tissue cell-type specific markers and co-expression to query how well neural organoids recapitulate primary tissue cell-type specific biology. We sample primary brain tissue data over the first and second trimesters and across 15 different developmentally defined brain regions, extracting lists of cell-type markers that define broad primary tissue cell-type identity regardless of temporal, regional, or technical variation (Fig. 1A). We derive co-expression networks from individual primary tissue and organoid datasets as well as aggregate co-expression networks across datasets (Fig. 1C). From these networks, we assess the strength of co-expression within primary tissue cell-type marker sets as well as the preservation of co-expression patterns between primary tissue and organoid data (Fig. 1D-E). We also provide an R package to download our primary tissue reference co-expression network to assay new neural organoid data using simple, meaningful, and fast statistics (Fig. 1F). By constructing robust primary tissue cell-type representations through meta-analytic approaches, we demonstrate the preservation of primary tissue cell-type co-expression provides both specific and generalizable characterization of the primary tissue fidelity of human neural organoids.

**Figure 1.**
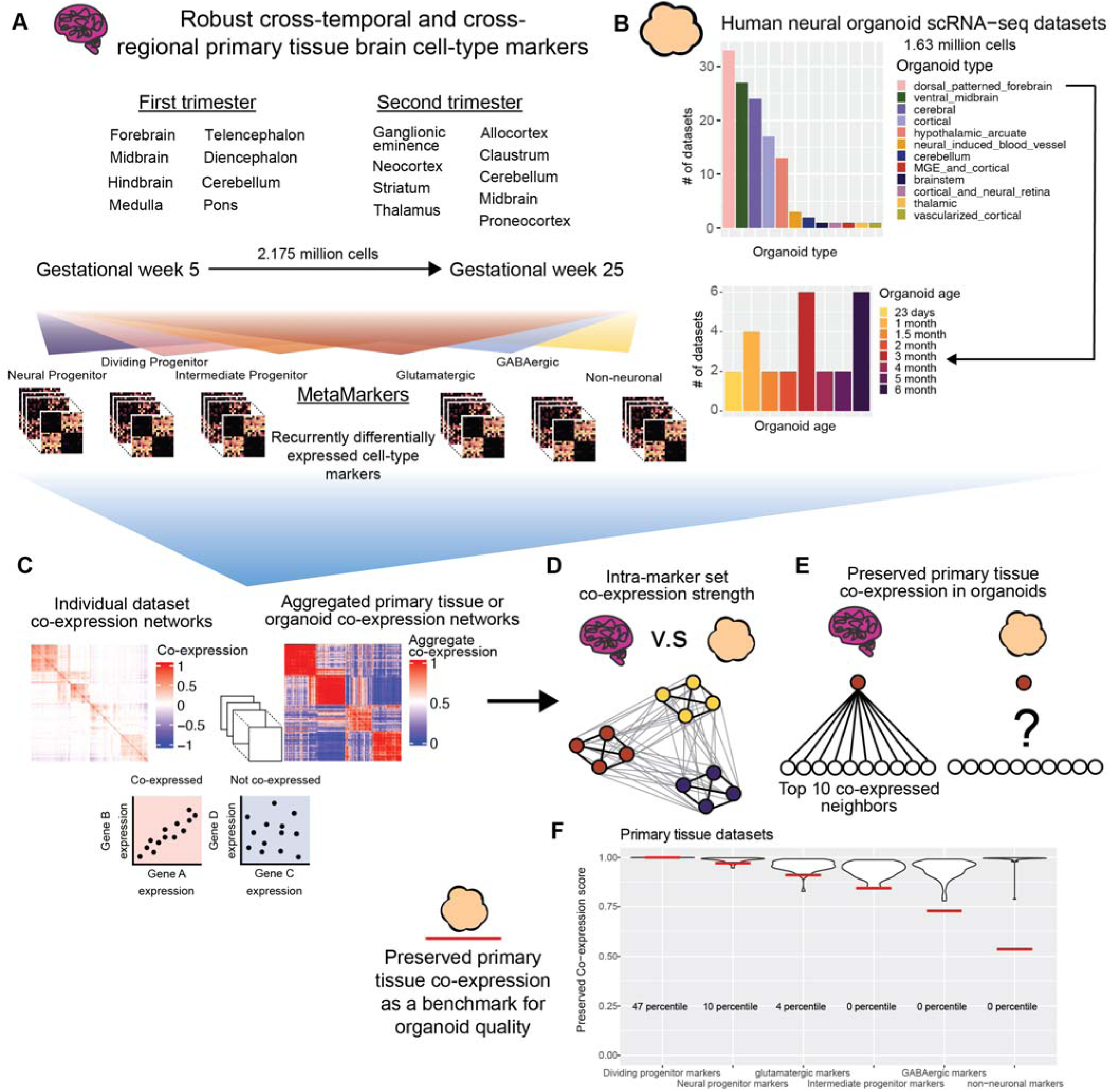
Using meta-analysis to quantify preserved primary tissue co-expression in organoids. **A** Collection of annotated primary tissue brain scRNA-seq datasets, ranging from gestational week (GW) 5 to 25 and sampling from 15 developmentally defined brain regions. The primary tissue datasets are annotated at broad cell-type levels (Neural Progenitor, Dividing Progenitor, Intermediate Progenitor, Glutamatergic, GABAergic, and Non-neuronal) and these annotations are used to compute MetaMarkers, cell-type markers identified through recurrent differential expression. **B** Collection of human neural organoid scRNA-seq datasets, sampling from 12 different differentiation protocols. Included is an annotated temporal forebrain organoid dataset. **C** Example of a sparse co-expression network derived from a scRNA-seq data and an example of an aggregate co-expression network averaged over many scRNA-seq datasets. The aggregate network enhances the sparse signal from the individual network. **D** Schematic showing a quantification of intra-marker set co-expression **E** Schematic showing a quantification for the strength of preserved co-expression between two co-expression networks, measuring the replication of the top 10 co-expressed partners of an individual gene across the networks. **F** Example plot from the preservedCoexp R library, placing cell-type specific preserved co-expression scores of an example forebrain organoid dataset in reference to scores derived from primary tissue datasets. Red lines denote the percentile of the organoid cell-type scores within the primary tissue distributions.

## Results

### Meta-analytic framework for primary tissue/organoid comparisons

We reason that, if they exist, primary tissue cell-type specific signals robust to temporal, regional, and technical variation will constitute *in vivo* standards applicable to any organoid dataset regardless of time in culture or differentiation protocol. We first show it is possible to learn sets of marker genes that define broad primary tissue cell-types (Fig. 1A, Supp. Table 1) across timepoints (gestational weeks GW5-GW25) and brain regions (15 developmentally defined brain regions) through a meta-analytic differential expression framework (Fig. 1A, Fig. 2A-B). We then compare co-expression within these marker sets between primary tissue and organoid data to quantify the degree organoids preserve primary tissue cell-type specific co-expression. An important aspect of our analysis is our cross-validation of primary tissue differential expression and co-expression. We employ a leave-one-out cross-validation approach when learning robust differentially expressed marker genes from our annotated primary tissue datasets (2,174,934 cells, 37 datasets) and we interrogate co-expression of our primary tissue marker genes within a large cohort of unannotated primary tissue datasets (776,343 cells, 14 datasets). This approach ensures we are extracting primary tissue markers and co-expression relationships independent of temporal, regional, and technical variation, a powerful approach for deriving broad primary tissue signatures appropriate for comparison to a wide range of organoid datasets.

**Figure 2.**
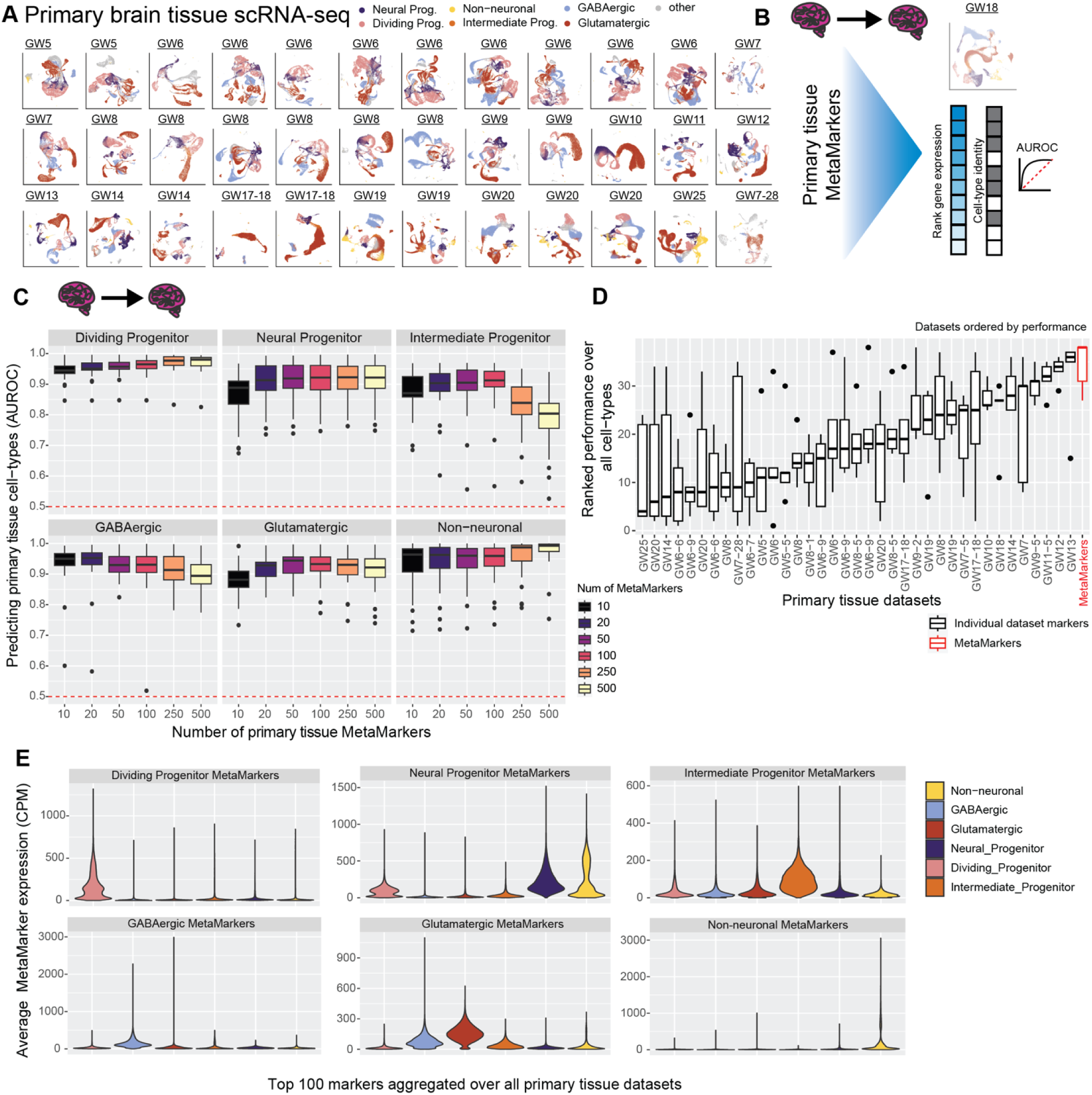
Meta-analytic primary tissue cell-type markers. **A** Annotated UMAPs of the annotated primary tissue brain scRNA-seq datasets. **B** Example of our leave-one-out cross-validation approach for learning primary tissue MetaMarkers and testing the markers’ capacity for predicting annotations in the left-out dataset, quantified with the AUROC statistic. **C** Meta-analytic primary tissue markers have high performance in predicting primary tissue cell-type annotations. Boxplot distributions of the AUROC statistic for predicting cell-type annotations across all leave-one-out combinations of our annotated primary tissue datasets, with an increasing number of MetaMarkers used for predicting cell-type annotations on the x-axis. **D** MetaMarkers have the highest performance in predicting primary tissue cell-type annotations. Boxplots of marker gene-set performances. Gene-sets are the top 100 cell-type markers from individual primary tissue datasets compared to the MetaMarker performance. Performances for each cell-type in individual primary tissue datasets are presented in Supp. Fig. 1A. Datasets are ordered by their median performance. **E** Averaged distributions of gene expression for the top 100 MetaMarkers demonstrating clear cell-type specificity. This is performed with a leave-one-out cross-validation, with individual dataset distributions reported in Supp. Fig. 1B.

### Cross-temporal and -regional primary tissue cell-type markers

To learn markers that define broad primary tissue cell-types (see methods), we apply the MetaMarkers^44^ framework to our cross-temporal and -regional annotated primary tissue datasets (Fig. 2A-B). MetaMarkers uses robust differential expression statistic thresholds (log2 fold-change >= 4 and FDR-adjusted p-value <= 0.05) for determining whether a gene is differentially expressed (DE) within individual datasets, then ranks all genes via the strength of their recurrent DE across datasets (see methods). We test the generalizability of our primary tissue MetaMarker gene sets in predicting primary cell-types by employing a leave-one-out primary tissue cross-validation (Fig. 2A-B). We construct an aggregate expression predictor in the left-out dataset using MetaMarkers learned from the remaining datasets (see methods), quantifying how well the MetaMarker gene sets predict the left-out cell-type annotations with the area-under-the-receiver-operating-characteristic curve statistic (AUROC, Fig. 2B-C). The AUROC is the probability of correctly prioritizing a true positive (e.g., cell of the right type) above a negative, (e.g., cell of the wrong type), given some predictor of the positive class, in this case, aggregate cell-type marker expression.

Starting with just the top 10 primary tissue MetaMarkers per cell-type, we achieve a mean AUROC across all primary tissue datasets of 0.944 ± 0.0280 SD, 0.864 ± 0.0796 SD, 0.873 ± 0.0676 SD, 0.937 ± 0.0669 SD, 0.879 ± 0.0535 SD, and 0.931 ± 0.0737 SD, for dividing progenitors, neural progenitors, intermediate progenitors, GABAergic neurons, glutamatergic neurons, and non-neuronal cell-types respectively (Fig. 2C). These extremely high performances demonstrate that even a small number of meta-analytically derived primary tissue cell-type markers have high utility in predicting primary tissue cell-type annotations regardless of temporal and regional variability. For all following analysis, we take the top 100 MetaMarkers per cell-type as robust representations of our 6 broad primary tissue cell-type annotations (average AUROC >= 0.90 except for intermediate progenitors: 0.897 ± 0.0777 SD), with the 100 MetaMarkers achieving modest increases in performance over the top 10 MetaMarkers for all cell-types except GABAergic cells (Fig. 2C, mean AUROC for 100 GABAergic MetaMarkers: 0.922 ± 0.0777 SD). When comparing MetaMarkers to markers derived from individual primary tissue datasets, we find the MetaMarkers are consistently top performers in predicting primary tissue annotations (Fig. 2D), with MetaMarkers producing the top results for intermediate progenitors, glutamatergic neurons, and GABAergic neurons (Supp. Fig. 1), as well as comparable performance to top individual datasets for dividing progenitors, neural progenitors, and non-neuronal cell-types (Supp. Fig. 1).

We explore the primary tissue MetaMarker sets further by computing the average expression of the top 100 MetaMarkers for our 6 annotated cell-types across all cells within our 37 annotated primary tissue datasets (Fig. 2E), continuing our leave-one-out approach. Each annotated primary tissue cell-type expresses the corresponding matched MetaMarker set over all other MetaMarker sets, with the exception of some off-target expression for the neural progenitor MetaMarkers in astrocytes (aggregated over all datasets Fig. 2E, individual datasets Supp. Fig. 1B). This demonstrates our MetaMarker gene sets act as robust cell-type markers in aggregate across all first and second trimester timepoints (Fig. 2E, Supp. Fig. 1B). Additionally, we investigate the expression of the top 100 MetaMarker gene sets across annotated primary brain regions, demonstrating each primary tissue cell-type maximally expresses the corresponding primary tissue MetaMarker set across all annotated brain regions (Supp. Fig. 2A-B). Overall, we are able to meta-analytically extract cell-type markers that define broad primary tissue cell-types independent of temporal and regional variation.

### Broad primary tissue cell-type markers capture organoid temporal variation

After extracting meta-analytic cell-type markers that capture broad primary tissue temporal and regional variation, we can test how well these markers also capture organoid temporal and regional (protocol) variation. We start with a large-scale temporal organoid atlas^38^ derived from a forebrain differentiation protocol containing timepoints ranging from 23 days to 6 months in culture. When comparing primary tissue and organoid data along a temporal axis, one might expect younger primary tissue expression data to be a better reference for younger organoid cell-types (better able to predict cell-types) and vice-versa for older primary and organoid data (Supp. Fig. 3A). We test this relationship using the same AUROC quantification as in Figure 1C, but now using the top 100 primary tissue cell-type markers per primary tissue dataset to predict organoid cell-type annotations across all organoid timepoints (Supp. Fig. 3B, see methods).

We observe highly consistent performance across all primary tissue datasets (GW5 – GW25) when predicting organoid cell-types regardless of the organoid timepoint (Supp. Fig. 3B). The average difference in AUROC scores when predicting organoid cell-types using either our youngest (GW5) or oldest (GW25) primary data is 0.000382 ± 0.0357 SD, 0.141 ± 0.192 SD, 0.139 ± 0.0317 SD, 0.00171 ± 0.113 SD and 0.119 ± 0.216 SD for dividing progenitors, neural progenitors, glutamatergic neurons GABAergic neurons, and non-neuronal cells respectively (No annotated intermediate progenitors in the GW25 primary tissue dataset). This demonstrates strikingly consistent performance across distant primary tissue timepoints, highlighting that broad primary tissue cell-type signatures are applicable as reference for organoid cell-types regardless of the primary tissue or organoid timepoint. The one exception is for neural progenitors, where there seemingly is a temporal shift in performance with younger primary tissue datasets predicting younger organoid annotations over older organoid annotations and vice-versa for older primary tissue/organoid data (Supp. Fig. 3B). However, a subset of the young GW6-8 primary tissue datasets report sharp increases in performance predicting older organoid timepoints in opposition to other GW6-8 primary tissue datasets, suggesting variance in performance is driven by intersections between the quality of individual organoid and primary tissue datasets rather than overarching temporal variability. Importantly, our lists of top 100 primary tissue MetaMarkers perform comparably to marker sets from individual primary tissue datasets, with less variance in performance across the organoid timepoints for the differentiated cell-types (mean AUROC variance across organoid timepoints for individual primary tissue datasets vs. primary MetaMarker variance; glutamatergic: 0.0147, 0.00672, GABAergic: 0.00487, 0.00201, non-neuronal: 0.00733, 0.00647, Supp. Fig. 3B). This demonstrates our meta-analytic primary tissue cell-type markers robustly capture organoid temporal variation.

### Broad primary tissue cell-type markers capture organoid protocol variation

We assess whether our primary tissue MetaMarker gene sets capture organoid variation outside the annotated forebrain temporal organoid atlas by performing principal-component analysis (PCA) across all organoid datasets, representing data from 12 different differentiation protocols. Our lists of 100 primary tissue MetaMarkers are consistently heavily weighted in the first PC across organoid datasets (Supp. Fig. 3C-D). While a large portion of PC1-weighted genes are dividing progenitor MetaMarkers (representing cell-cycle signal), markers for non-dividing fetal cell-types also comprise those genes consistently heavily weighted in PC1 across organoid datasets (Supp. Fig. 3C-D).

### Aggregate organoid co-expression weakly captures primary tissue co-expression

Our primary tissue MetaMarkers that capture both primary tissue and organoid temporal/regional variation enable assessments of cell-type specific co-expression between arbitrary primary tissue and organoid datasets. One normally would need matched cell-type annotations across datasets to compare cell-type specific biology, but here we couple our meta-analytically derived cell-type markers with gene co-expression quantifications, which do not rely on cell-type annotations, to extract cell-type specific co-expression from any given scRNA-seq dataset. Practically, if organoids are producing cell-types functionally identical to primary tissue cell-types, we would expect near identical co-expression relationships within our primary tissue MetaMarker gene sets across primary tissue and organoid datasets.

We first explore marker set co-expression within our unannotated primary tissue datasets, which were not included in deriving our primary tissue MetaMarker sets. The aggregate (Fig. 3A, see methods) unannotated primary tissue co-expression network nearly perfectly constructs cell-type specific co-expression modules when hierarchically clustering the co-expression of our top 100 primary tissue MetaMarker gene sets (Fig. 3B). Turning to the aggregate organoid co-expression network, while some cell-type co-expression structure exists, it is much weaker than the unannotated primary tissue co-expression with less well-defined intra-gene set co-expression relationships (Fig. 3B). We quantify this through the Adjusted Rands Index (ARI) metric, comparing the MetaMarker clustering through co-expression in any given network to the perfect clustering of MetaMarker gene sets by cell-type. We perform this quantification for both the aggregate co-expression networks (diamond, triangle, and square special characters, Supp. Fig. 4A) and for all individual primary tissue and organoid co-expression networks (boxplots, Supp. Fig. 4A). Individual organoid networks perform worse than individual primary tissue networks on average, with the aggregate organoid network additionally underperforming compared to the aggregate primary tissue networks, though within the range of individual primary tissue networks (Supp. Fig. 4A). In aggregate, organoid co-expression weakly captures broad primary tissue cell-type specific co-expression. This is potentially explained through the directed nature of the vast majority of organoid datasets we investigate, which may more accurately produce particular lineages (excitatory or inhibitory neurons as an example) rather than the comprehensive cell-types/lineages present within primary tissue data. We explore cell-type specific co-expression within individual datasets further in the following analysis.

**Figure 3.**
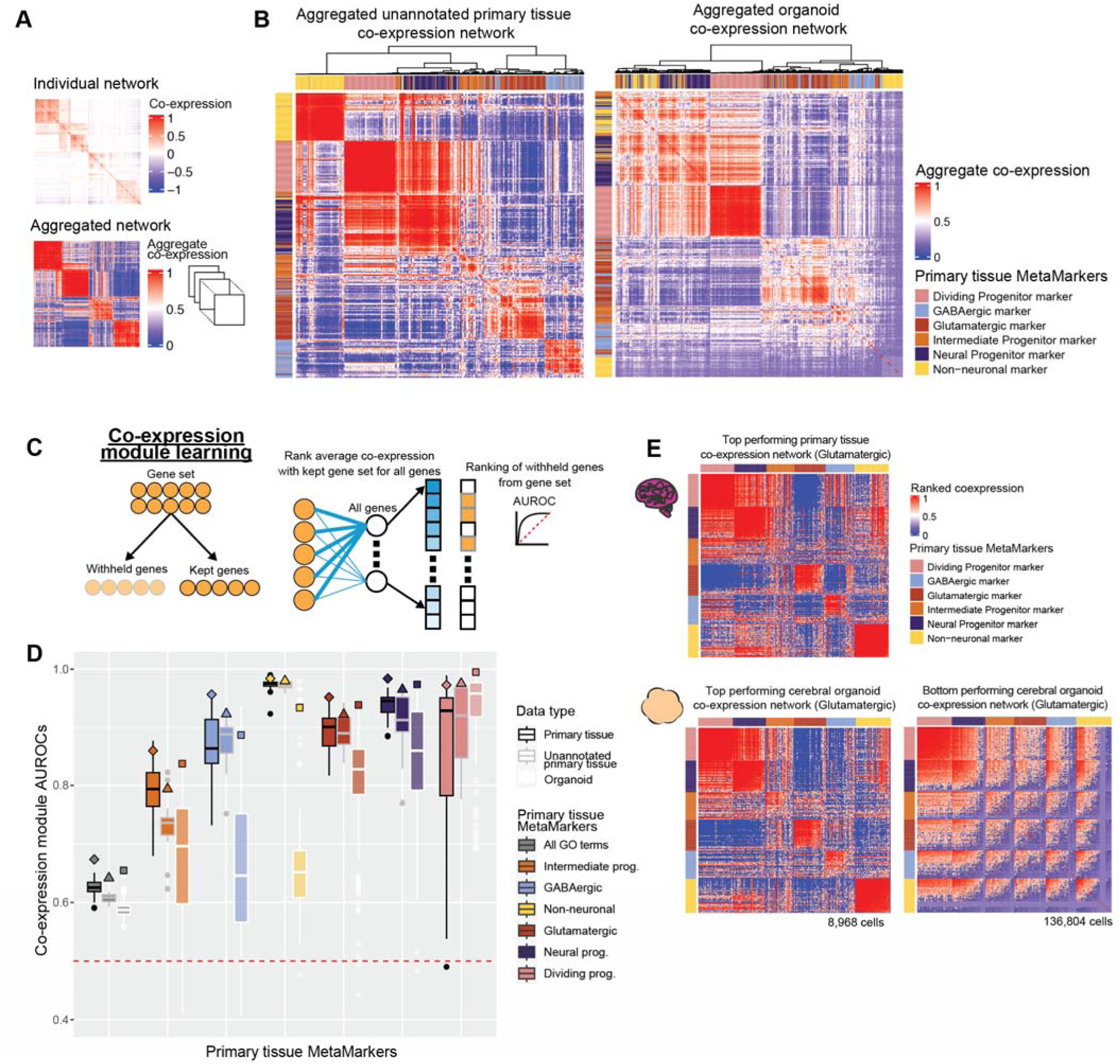
Neural organoids vary in recapitulating primary tissue cell-type marker set co-expression. **A** Example of a sparse co-expression network derived from a scRNA-seq data and an example of an aggregate co-expression network averaged over many scRNA-seq datasets. The aggregate network enhances the sparse signal from the individual network. **B** Marker gene-sets show clear cell-type clusters via their co-expression relationships in primary tissue and organoid networks. The aggregated co-expression networks for the unannotated primary tissue datasets and organoid datasets, showing the hierarchically clustered co-expression of the primary tissue MetaMarkers for the 6 cell-types. **C** Schematic for the co-expression module learning framework, measuring the co-expression strength within an arbitrary gene-set compared to the rest of the genome, quantified with the AUROC statistic. **D** Distributions of co-expression module AUROCs for individual annotated primary tissue, unannotated primary tissue, and organoid datasets for the co-expression strength of the MetaMarker gene-sets for the 6 cell-types. The grey ‘All GO terms’ distributions report the average co-expression module AUROC across all GO terms for each individual dataset. Co-expression module AUROCs for the aggregate co-expression networks are denoted with the special characters. **E** Top primary tissue and top and bottom organoid co-expression networks based on Glutamatergic co-expression module AUROCs. Genes are ordered within each MetaMarker gene set by their average intra-gene set co-expression.

### Organoid datasets vary in primary tissue cell-type marker set co-expression

Having broadly assessed co-expression across our MetaMarker gene sets, we then asked how well do organoids recapitulate primary tissue co-expression within each cell-type specific MetaMarker gene set. We score intra-gene set co-expression strength through a simple machine learning framework^45,46^, which quantifies whether genes in a given set are more strongly co-expressed with each other compared to the rest of the genome (Fig. 3C).

Co-expression module scores across the annotated and unannotated primary tissue datasets are largely comparable with the exception of a sharp decrease in intermediate progenitor performance for the unannotated primary tissue datasets (Fig. 3D). Six out of the fourteen unannotated datasets are sampled from either the ganglionic eminences or the hypothalamus, potentially explaining this decrease in performance and suggesting our intermediate progenitor MetaMarkers are enriched for signal from cortical areas. In contrast, performance is much more variable across the individual organoid datasets for all cell-types except the dividing progenitors, ranging from no signal (AUROC <= 0.50) to comparable results with primary tissue networks (Fig. 3D).

Importantly, the variation among organoid datasets for co-expression of the differentiated cell-types is likely influenced via the compositional variation in cell-types produced across directed and undirected differentiation protocols. A protocol that aims to produce a directed excitatory lineage organoid is not expected to produce inhibitory cell-types and thus should not necessarily exhibit strong inhibitory neuron co-expression. That is indeed the case when comparing our co-expression module scores by organoid protocols (Supp. Fig. 4B). For example, the dorsal patterned forebrain organoid protocol produces stronger excitatory module co-expression compared to inhibitory module co-expression.

In contrast, undirected organoid protocols are expected to produce a variety of lineages/cell-types comparable to those present in primary tissue samples and should exhibit consistent strong co-expression across cell-types. Instead, we report the undirected organoid protocols (cortical, cerebral) as the more variable protocols for producing strong cell-type co-expression (Supp. Fig. 4B). Visualizing the top and bottom performing cerebral organoid co-expression networks for glutamatergic co-expression reveals the extent of this variability, in comparison to the top performing primary tissue co-expression network (Fig. 3E). The top performing organoid network produces near identical intra- and inter-cell type co-expression relationships to the primary tissue dataset (Fig. 3E). Contrastingly, the bottom performing organoid co-expression network exhibits extensive off-target inter-cell type co-expression and extremely poor intra-cell type co-expression, essentially failing to recapitulate primary tissue cell-type co-expression (Fig. 3E). While variability in co-expression performance may reflect compositional differences among directed organoids, the dramatic range in performance across the undirected organoid datasets reveals extensive variability in fidelity to primary tissue.

### Organoid datasets vary in preserving gene-level primary tissue co-expression

We take our primary tissue/organoid co-expression comparisons a step further and ask how well individual organoid datasets preserve gene-level primary tissue co-expression relationships. For any given individual gene, we quantify whether that gene’s top co-expressed partners are preserved in one co-expression network compared to another (Fig. 4A). We use the aggregate co-expression network from the annotated primary tissue datasets as our reference co-expression network and test how well individual co-expression networks, either primary tissue or organoid, perform in preserving primary tissue gene-level co-expression patterns (Fig. 4A, top 10 co-expressed neighbors). We start by quantifying the preserved co-expression of genes within our primary tissue MetaMarker gene sets, using the average preserved co-expression AUROC as a measure of preserved co-expression for any given gene set (Fig. 4A). Across our 6 annotated primary tissue cell-types, primary tissue co-expression networks deliver consistently high performance for preserved co-expression scores of our primary tissue MetaMarker gene sets (Fig. 4B, mean preserved co-expression score across cell-types and primary tissue datasets: annotated 0.970 ± 0.0241 SD, unannotated 0.963 ± 0.00940 SD). This indicates that across the highly temporally and regionally diverse primary tissue data, the co-expression relationships of our MetaMarker gene sets are incredibly highly preserved, again reflecting the temporally and regionally robust nature of our primary tissue cell-type markers.

**Figure 4.**
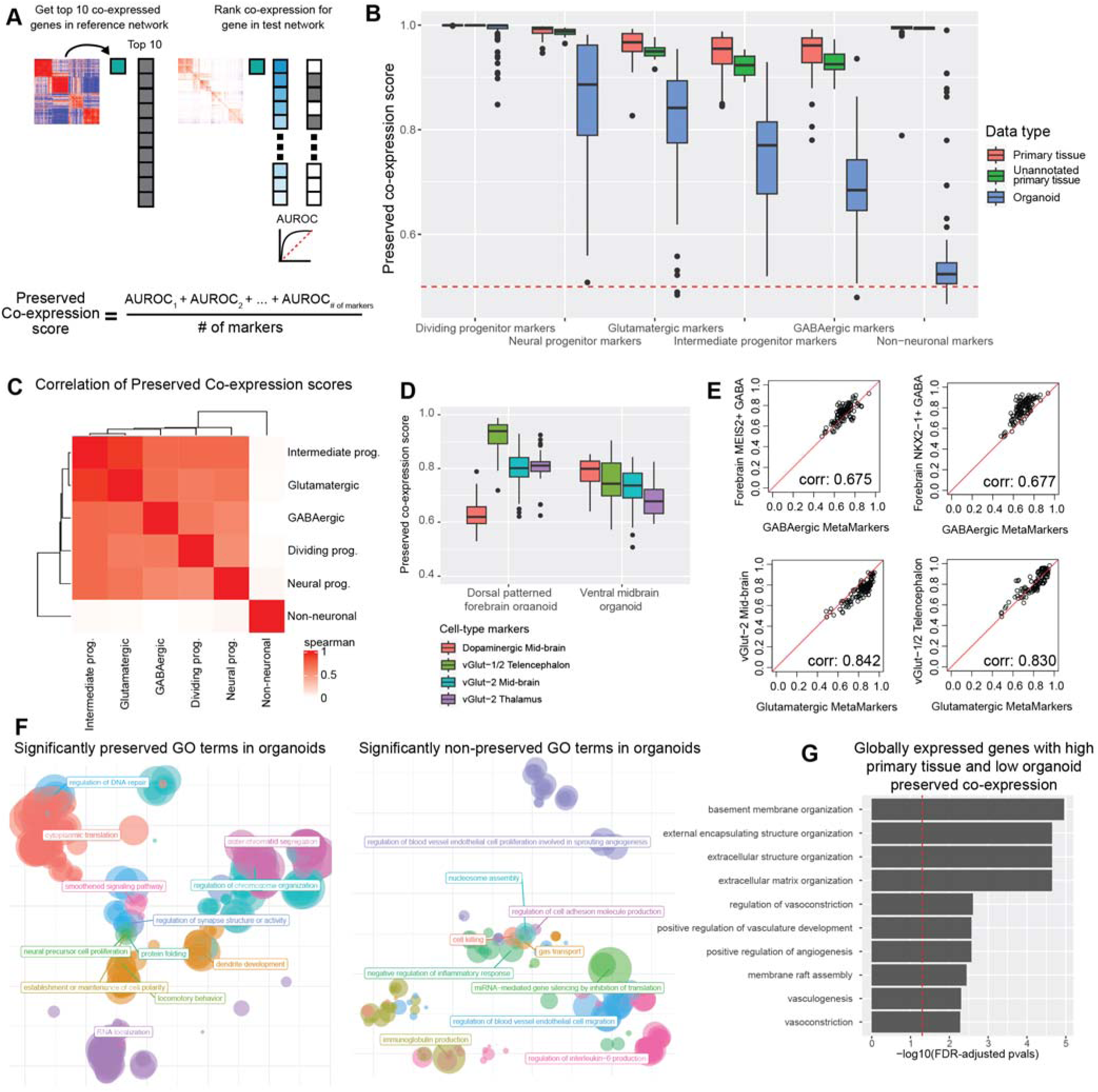
Neural organoids vary in their preservation of primary tissue gene-level co-expression. **A** Schematic showing the quantification for gene-level preserved co-expression. The preserved co-expression score for any given gene-set is the average preserved co-expression AUROC across all genes within that gene set. **B** Organoids strongly vary in preserved primary tissue cell-type specific co-expression in comparison to fetal data. Boxplot distributions show the preserved co-expression scores for the primary tissue MetaMarker gene-sets of the 6 cell-type annotations across all individual networks. **C** The majority of cell-types are significantly correlated in preserved co-expression within organoid networks. Spearman correlation matrix for the preserved co-expression scores for all 6 cell-type annotations across all individual organoid datasets. **D** Preserved co-expression scores computed from the dorsal patterned forebrain and ventral midbrain organoid datasets for the top 10 cell-type markers of various neural cell-types. **E** Scatter plots comparing the preserved co-expression score of the top 100 MetaMarkers against the top 10 markers (no overlaps in gene sets) for various neural cell-types per organoid dataset. Spearman correlation coefficients are reported in the bottom right corner. **F** Scatter plots summarizing the semantic distances of GO terms that are significantly preserved or non-preserved between the aggregate annotated primary tissue and organoid co-expression networks. **G** Organoids globally fail to preserve primary tissue co-expression of ECM and vascular related genes. Bar plot detailing the top 10 GO terms from a GO enrichment test of the 76 genes with high and low preserved co-expression AUROCs within primary tissue networks and organoid networks respectively. The preserved co-expression for each individual gene from primary tissue networks and organoid networks is reported in Supp. Fig. 6B.

In contrast, individual organoid datasets vary substantially in preserved co-expression scores across our primary tissue MetaMarker gene sets (Fig. 4B). As before with our quantification of intra-gene set co-expression, compositional variation across directed organoid protocols may influence the variation in performance for differentiated cell-types, especially for excitatory and inhibitory neurons (Supp. Fig. 5). To explore the effects of likely compositional variation, we compared preserved co-expression scores of organoids grown in a vertical shaker versus an orbital shaker^47^, where the original authors reported either a GABAergic or glutamatergic character for organoids grown in either a vertical or orbital shaker respectively. We report that organoids grown in an orbital shaker produce higher preserved primary tissue co-expression scores for intermediate progenitors and glutamatergic cell-types whereas organoids grown in a vertical shaker produce higher scores for GABAergic cell-types, in agreement with the authors original observations (3 replicates each, glutamatergic, intermediate progenitor, GABAergic; Orbital: 0.896 ± 0.0105 SD, 0.795 ± 0.00146 SD, 0.665 ± 0.0302 SD. Vertical: 0.644 ± 0.0126 SD, 0.686 ± 0.0167 SD, 0.762 ± 0.00589 SD). However, regardless of putative compositional variation across organoid protocols and/or treatments, we demonstrate that organoids from all sampled protocols consistently fail to preserve glutamatergic or GABAergic primary tissue co-expression at a level comparable to primary tissue (Supp. Fig. 5, Preserved Co-expression score ∼0.90 or higher). This suggests a persistent remaining biological gap in the fidelity of organoid neurons in reference to primary tissue neurons. Organoids across protocols additionally exhibit near-zero preservation of non-neuronal primary tissue co-expression, suggesting organoids generally do not produce or produce extremely dysregulated non-neuronal cell-types (Fig. 4B, Supp. Fig. 5).

Additionally, we report extensive variability in the preservation of primary tissue progenitor co-expression across organoid datasets. Again, regardless of the differentiation protocol, organoids persistently fail to achieve comparable preserved co-expression of the neural and intermediate progenitor MetaMarkers in reference to primary tissue data (Fig. 4B, Supp. Fig. 5). In contrast to the differentiated cell-types, where compositional variation of lineages across differentiation protocols may explain performance, progenitors are present within every neural organoid. Variability in preserved co-expression of progenitor Meta-Markers is likely a stronger reflection of the fidelity to primary tissue of any given organoid dataset. Interestingly, a vascularized organoid protocol produces the highest preserved co-expression of the neural and intermediate progenitors as well as the glutamatergic and GABAergic cell-types. This suggests that vascularized organoids are particularly adept at producing cell-types with high fidelity to primary tissue, but also that the preservation of co-expression is associated across cell-types. We quantify this by computing correlations of preserved co-expression scores between the 6 MetaMarker gene sets across all organoid datasets and find significantly positive correlations (FDR-adjusted p-value < .001) across all comparisons with the exception of the non-neuronal cell-type (Fig. 4C, non-neuronal FDR-adjusted p-values range from 0.650 to 0.731). This indicates preserved primary tissue co-expression is a global feature of organoid datasets. For example, if an organoid is producing neural progenitors that preserve primary tissue co-expression, that organoid is likely producing other cell-types that preserve primary tissue co-expression.

Neural organoids are commonly employed for the study of diverse disease mechanisms through various perturbations. We tested the relevance of our preserved co-expression scores for quantifying primary tissue fidelity across normal and perturbed organoids. A subset of our organoid datasets come from studies that performed diverse perturbations (22q11.2 deletion, SMARCB1 knockdown, exposure to Alzheimer’s serum, SETBP1 point mutations, amyotrophic lateral sclerosis patient-derived organoids). We compare the MetaMarker preserved co-expression scores between normal and perturbed organoids and find only a single significant difference across all cell-type MetaMarker sets (Intermediate Progenitor normal vs. mutant preserved co-expression score FDR-adjusted p-value: 0.0295, Supp. Fig. 6A). This demonstrates our broad primary tissue cell-type co-expression signatures are also applicable for comparison with organoids in perturbation experiments.

### Fidelity of finer resolution cell-types through preserved co-expression

While our broad cell-type annotations are useful for unifying meta-analysis across heterogeneous primary tissue and organoid datasets, it is also of interest the degree neural organoids are capable of producing primary tissue cell-types at a finer resolution. As our approach for quantifying the preservation of co-expression is derived from a genome-wide co-expression network of primary neural tissue, we can also putatively assess preserved co-expression of more specific cell-type markers. We investigate preserved co-expression of more specific cell-type markers by utilizing marker genes derived from a morphogen screen in neural organoids that reported the production of extensive neural cell-type diversity^48^. As examples of protocol specific trends, we show the dorsal patterned forebrain organoid preserves co-expression of telencephalic excitatory neuron markers over markers for mib-brain and thalamic excitatory neurons as well as dopaminergic mid-brain neurons (Fig. 4D). Similarly, the ventral mid-brain organoid protocol, which reported production of dopaminergic neurons, preserves co-expression of dopaminergic neuron markers over excitatory neuron markers on average (Fig. 4D). Extending across all the organoid datasets, we demonstrate preserved co-expression of fine resolution cell-types exhibit high correlations with the preservation of our broader class-level markers for several Glutamatergic and GABAergic cell-types (Fig. 4E). In summary, our results show that disruption of co-expression at one level of cell-type hierarchy captures disruption at finer levels, suggesting a single score for organoid fidelity can capture shared variation. More generally, our quantification for preserved co-expression in organoids can also be applied to the study of finer resolution cell-types to study variation from the shared baseline.

### Genome-wide preservation of co-expression reveal consistent organoid deficits

After revealing cell-type specific variation for preserving primary tissue co-expression within organoids, our co-expression networks additionally allow genome-wide assessments of preserved co-expression. We extend our analysis via GO terms to quantify preserved primary tissue co-expression within organoids across the whole genome. GO terms with significantly preserved primary tissue co-expression (see methods) in organoids are mostly related to basic cellular functions like response to DNA damage and protein translation, as well as GO terms related to neurodevelopment (Fig. 4F). GO terms that significantly lack preservation of primary tissue co-expression are largely related to angiogenesis or immune function (Fig. 4F), concordant with the fact that organoids lack vasculature and an immune system. These results demonstrate quantifications of preserved co-expression can capture known biological deficits in neural organoids.

While GO terms are useful for partitioning the genome into functional units for comparison, our co-expression networks also enable assessments of preserved co-expression for individual genes. As a particular use-case, we search for genes with exceptionally high preserved primary tissue co-expression across primary tissue datasets that also have poor preserved primary tissue co-expression across organoid datasets. We only consider genes that have some measurable expression in every organoid and primary tissue dataset and compute the average preserved co-expression AUROC for each gene across the organoid and primary tissue datasets (Supp. Fig. 6B). The top 10 enriched GO terms for genes (76 in total) with high primary tissue (average AUROC >= 0.99) and low organoid (average AUROC < 0.70) preserved co-expression are related to extra-cellular matrix (ECM) and vascular characterizations (Fig. 4G). The poor conservation of genes related to vasculature can be explained by the absence of vascularization in the vast majority of our organoid datasets. The subset of these 76 genes in the ECM GO terms are CAV1, CAV2, COL4A1, CTSK, ENG, LAMB1, LAMC1, NID1, NID2, DDR2, and VWA1. Notably, these genes produce collagen and laminins, components of Matrigel, the artificial ECM typically included in organoid cultures. These results highlight preserved primary tissue co-expression of ECM-related genes as a particularly consistent deficit across neural organoids, suggesting that investigations into the signaling between artificial ECM and cells in organoid cultures may be a route forward for general improvements of organoid fidelity.

In summary, we interrogate co-expression in organoids at multiple levels, revealing organoids vary in preserving primary tissue co-expression at gene-, cell-type, and whole genome resolutions through the use of a robust aggregate primary tissue co-expression network. We demonstrate the applicability of our approach for quantifying primary tissue fidelity in organoids against a variety of use-cases, such as comparing different culture conditions (vertical vs orbital shaking), comparing normal and perturbed organoids, and investigating preserved co-expression of individual genes and fine resolution cell-type markers.

### Temporal variation in organoid preservation of primary tissue co-expression

We score preserved co-expression in organoids using the aggregate primary tissue co-expression network (GW5-25), which by design aims to capture signal robust to temporal variation. To investigate temporal trends in organoid co-expression, we employ a similar approach as when predicting organoid cell-type annotations (Supp. Fig. 3), this time quantifying the preservation of primary tissue co-expression for the top 100 cell-type markers per individual primary tissue dataset across all organoid timepoints (Fig. 5A-B). We uncover a broad temporal shift in the preservation of primary tissue co-expression within organoids across all cell-types, with younger organoids (23 days – 1.5 months) as the top performers for mostly first trimester primary tissue co-expression transitioning to older organoids (2 – 6 months) as top performers for mostly second trimester primary tissue co-expression (Fig. 5B). This temporal shift is broadly consistent across the cell-types, beginning around GW9-10 (Fig. 5B). Our approach in predicting organoid annotations in Figure 2 is based on aggregate marker expression and did not produce temporally variable results, whereas our approach here comparing preserved co-expression of the same marker genes does produce temporally variable results. This indicates that the co-expression relationships of genes rather than their expression levels better capture temporal variation in developing systems.

**Figure 5.**
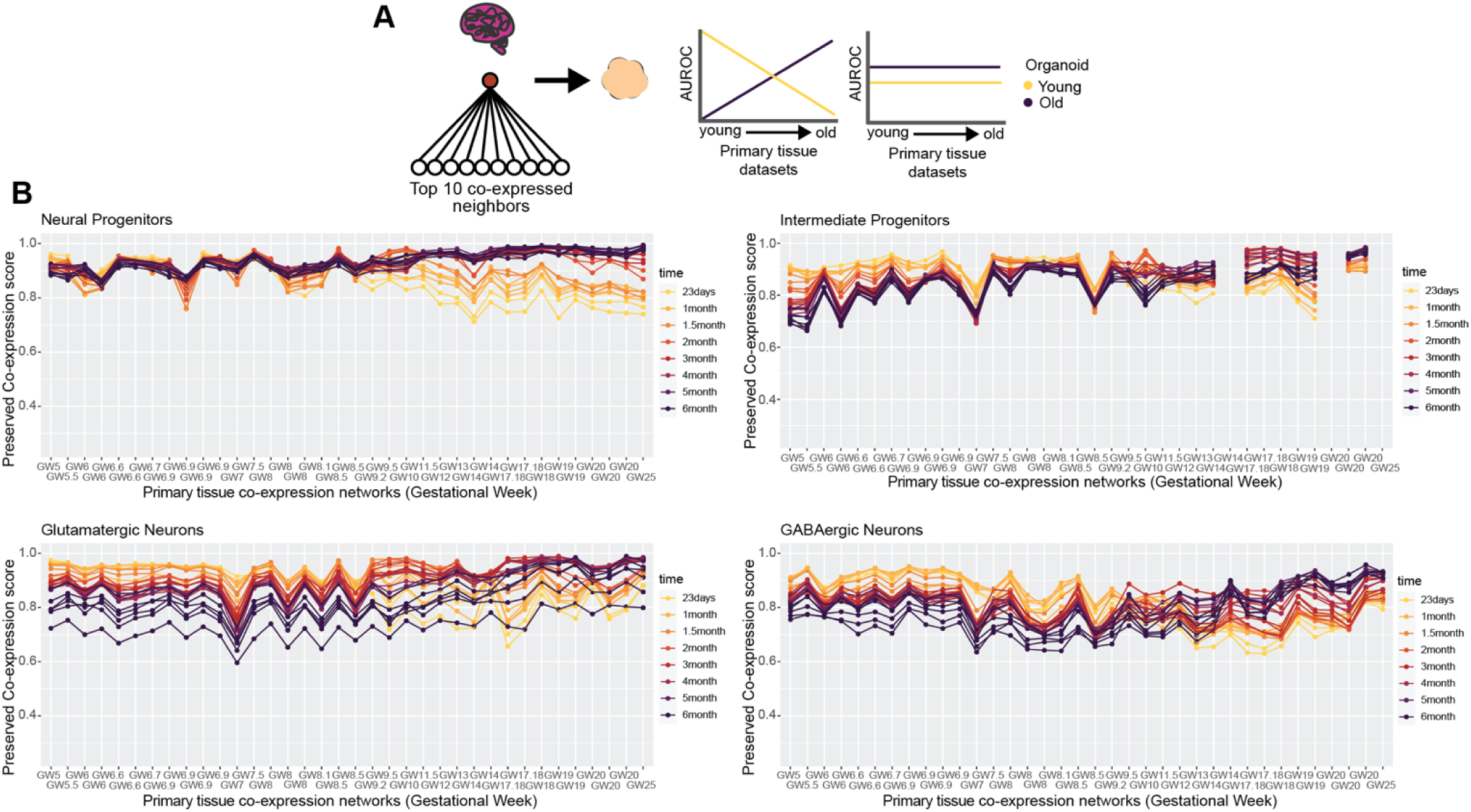
Neural organoids capture temporal dynamics in primary tissue co-expression. **A** Schematic showing two potential outcomes when comparing the preserved co-expression between primary tissue and organoid data on a temporal axis. There may be a temporal relationship, with younger organoids recapitulating younger primary tissue co-expression over older primary tissue co-expression and vice versa for older organoids, or there may be no temporal relationship. **B** Organoid co-expression models temporal trends in primary tissue co-expression. Line plots showing the preserved co-expression scores computed from individual organoid co-expression networks for cell-type markers of individual primary tissue datasets. Primary tissue datasets on the x-axis are ordered from youngest to oldest.

### Organoids preserve developing brain co-expression over adult brain co-expression

We demonstrate temporal variation in developing brain co-expression relationships is captured by organoids, but only from the single forebrain organoid protocol used in the temporal organoid atlas. In order to extend analysis across all our organoid datasets and assess broad temporal variation in co-expression, we next investigate the preserved co-expression within organoids of both developing and adult brain co-expression relationships.

We construct an aggregate adult co-expression network from a medial temporal gyrus scRNA-seq dataset^49^. We compare the preserved co-expression scores of organoids for either developing or adult glutamatergic, GABAergic, and non-neuronal cell-types. Organoids unanimously preserve developing brain co-expression over adult co-expression (Supp. Fig. 6C) for glutamatergic and GABAergic cell-types with equally poor performance for the non-neuronal cells, again suggesting organoids generally fail to produce non-neuronal cell-types. We extend this analysis genome-wide and place organoids in context between developing and adult data by computing the average preservation of co-expression AUROC across all genes for organoid, developing, and adult co-expression using the annotated primary developing brain tissue network as the reference. The adult co-expression network produces a global preserved developing brain co-expression score of 0.591, indicating very poor performance across the genome in preserving developing co-expression relationships (Supp. Fig. 6D). Organoids vary substantially in their global preservation of developing brain co-expression with some organoid datasets performing comparably to the adult data. This result is largely influenced by the number of cells present within individual organoid datasets (Supp. Fig. 6D, corr 0.647, p-value < .001), suggesting a cell-sampling limitation for uncovering developing brain co-expression within organoids. However, organoid datasets report more variable global preserved co-expression scores compared to down-sampled developing brain data (Supp. Fig. 6D), indicating a remaining gap between primary developing brain tissue and organoid data not explained through cell number sampling alone.

We further explore the applicability of our preserved co-expression quantifications for investigating temporal variation through a study that tested the limits of neuronal maturation in organoids. This study generated data from human cortical organoids either transplanted or not into developing rat brains to test the limits of maturation organoids can achieve *in vitro*^50^. We compare the preservation of developing and adult co-expression between these age-matched non-transplanted and transplanted human cortical organoids. We report that while the non-transplanted organoids preserve developing co-expression over adult for glutamatergic and GABAergic markers (Supp. Fig. 6E, non-transplanted glutamatergic and GABAergic mean developing brain AUROCs: 0.798 ± 0.0278 SD, 0.698 ± 0.0208 SD. Non-transplanted glutamatergic and GABAergic mean adult AUROCs: 0.672 ± 0.0234 SD, 0.585 ± 0.0291 SD), the transplanted organoids have increased preservation of adult co-expression for glutamatergic markers (Supp. Fig. 6E, transplanted glutamatergic mean developing brain AUROCs: 0.759 ± 0.00909 SD. Transplanted glutamatergic mean adult AUROCs: 0.850 ± 0.0332 SD). This indicates the transplanted human organoids are adopting adult human glutamatergic co-expression, concordant with the original authors’ conclusions of increased maturation in transplanted organoids. The transplanted organoids additionally report increased preservation of both developing and adult non-neuronal marker co-expression, in agreement with the original authors’ observations of oligodendrocytes within transplanted organoids. By recapitulating known maturation dynamics in organoid models, we demonstrate the broad applicability of preserved co-expression quantifications for investigating a range of biological phenomenon in neural organoids.

### Variability in organoid co-expression is driven by marker gene expression

We investigate the impact of various technical features in our analysis on our co-expression results by assessing their correlation with our co-expression module scores and preserved co-expression AUROCs, focusing on technical features like sequencing depth, number of cells, etc. An important technical consideration for our analysis is ensuring all datasets have an identical gene namespace for meaningful comparisons of expression data.

We fit all datasets to the GO gene universe, dropping gene annotations not in GO or zero-padding missing GO annotations in individual datasets. Excessive zero-padding of genes within our MetaMarker gene sets may artificially lower co-expression module scores or preserved co-expression scores, though we find this relationship to be relatively weak with little impact on score variance (Supp. Fig. 7, R^2^ for co-expression module scores and zero-padding: 0.00267, 0.0165, 0.126, 0.0261, 0.0354, 0.00451, R^2^ for preserved co-expression and zero-padding: 0.0665, 0.322, 0.151, 0.0307, 0.0411, 0.00203 for neural prog., dividing prog., intermediate prog., glutamatergic, GABAergic, and non-neuronal cell-types respectively). Sequencing depth is also similarly found to have little impact on our co-expression module scores or preserved co-expression scores (Supp. Fig. 7). Rather, the features strongly related to performance are the number of cells in a dataset and the strength of marker set expression (Supp. Fig. 7, range of significant positive (p-value < .05) correlations between marker set expression or cell number and co-expression module scores or preserved co-expression scores: 0.204 – 0.809).

### Preservation of primary tissue co-expression as a generalizable quality control metric

As a general summary, our approach for quantifying preserved primary tissue co-expression across numerous organoid protocols reveal the axes on which organoids lie for recapitulating primary tissue co-expression relationships at gene, cell-type, and whole-genome resolutions. These assessments provide powerful quality control information, identifying which genes and/or cell-types organoids can or cannot currently model on par with primary tissue data. We make our methods accessible through an R package to aid in future organoid studies and protocol development, providing means for rapidly constructing co-expression networks from scRNA-seq data (Fig. 6A) as well as querying preserved co-expression of users’ data with our aggregate primary tissue brain co-expression network (Fig. 6A). Additionally, we make the results of our meta-analysis across primary tissue and organoid datasets available for users to place their data in reference to a field-wide collection (Fig. 6B).

**Figure 6.**
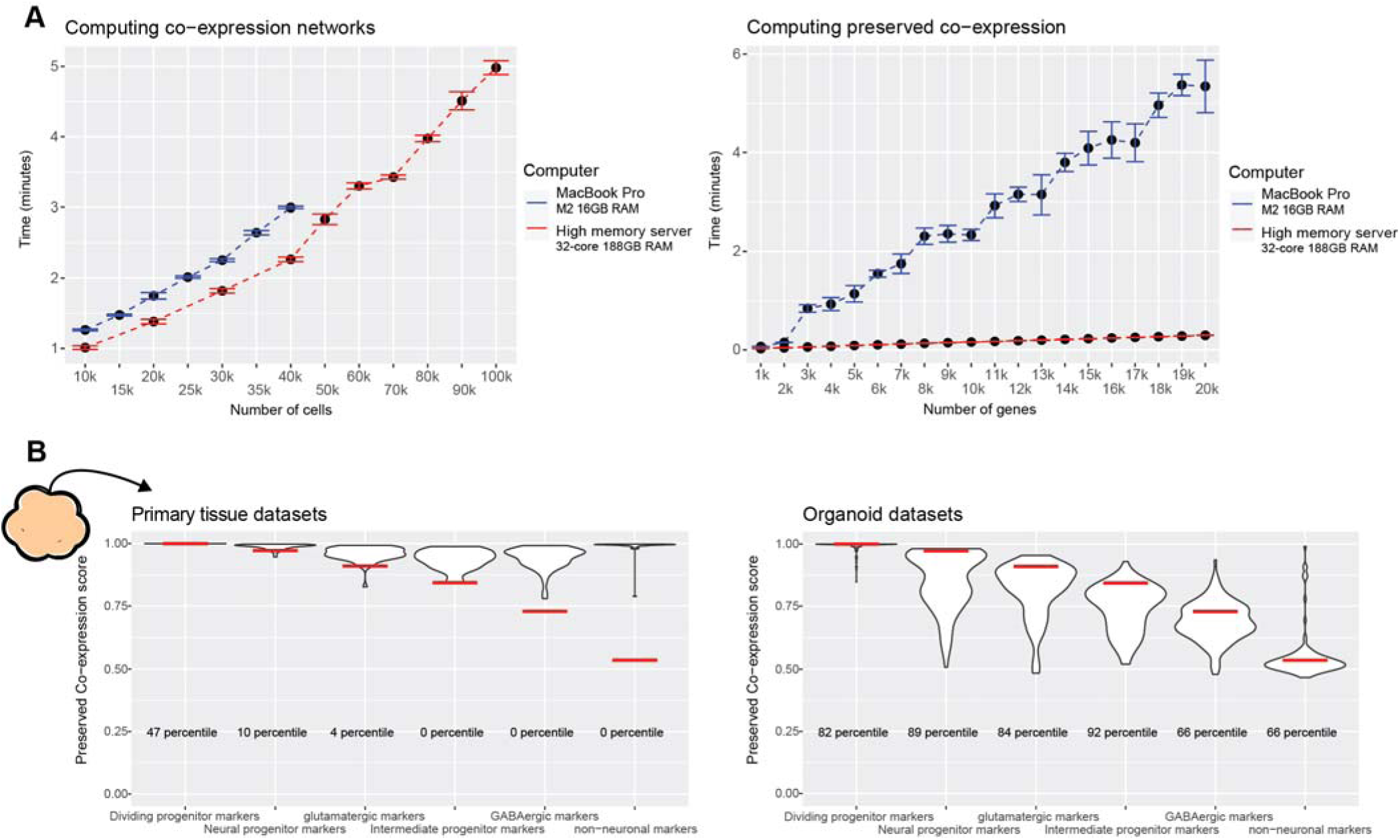
The preservedCoexp R package enables fast computation of preserved co-expression. **A** The preservedCoexp R package can compute co-expression networks and genome-wide preservation of co-expression in a few minutes even for low-memory computers. Line plots showing the computational time to either compute co-expression networks or preserved co-expression as the number of cells or genes increases. Points are the mean value from 10 replicates, with error bars depicting ± 1 standard deviation. **B** Example plot from the preservedCoexp R package, placing cell-type specific preserved co-expression scores of an example forebrain organoid dataset in reference to scores derived from primary tissue datasets or organoid datasets. Red lines denote the percentile of the forebrain organoid cell-type scores within either the primary tissue distributions or organoid distributions.

## Discussion

Through the use of meta-analytic differential expression and co-expression, we are able to provide cell-type specific measurements of human neural organoids’ current capacity to replicate primary tissue biology. We extracted broad cell-type markers that define primary brain tissue cell-types across a large temporal axis (GW5 – 25) and across numerous heterogenous brain regions to act as a generalizable primary tissue reference for organoids that also vary temporally and regionally (by protocol). By quantifying intra-marker set co-expression and the preservation of co-expression across networks, we revealed human neural organoids lie on a spectrum of near-zero to near-identical recapitulation of primary tissue cell-type specific co-expression in comparison to primary tissue data. We made our aggregate primary tissue reference data and methods for measuring preserved co-expression publicly available as an R package to aid in the quality control and protocol development of future human neural organoids.

Prior work comparing primary brain tissue and neural organoid systems demonstrated organoids can produce cell-types^11,12^ and morphological structures^27,43^ similar to primary tissues and are capable of modeling temporal^13,38,40^ and regional^3,12,28,29^ primary tissue variation. Multiple lines of evidence support these findings such as assessments of cytoarchitecture and cell-type proportions^3,11,16,23^, whole transcriptome and marker gene expression correlations^10,12^, and comparisons of co-expression modules^5,13,17,39^. Our meta-analytic approach is able to quantify these field-wide observations within a generalizable framework, recapitulating that organoids model broad primary tissue biology with our specific approach offering several key advancements for primary tissue/organoid comparisons. First, we derive quantifications of preserved primary tissue co-expression that can be extended from individual genes to the entire genome and, second, we place organoid co-expression in reference to robust meta-analytic primary tissue performance providing a general benchmark for protocol development and quality control across heterogeneous organoid systems.

Certainly, while comparisons between primary tissue and organoid systems at a high-resolution of cell-type annotation are of interest, our results centered on broad cell-types at the cell-class level constitute a critical foundation for these more fine-tuned investigations of organoids. Cell-type specification within the brain involves complex spatial and temporal mechanisms^51^ to produce the high cellular heterogeneity we observe, with the exact resolution of meaningful cell-type annotations still being actively debated and posing a general conceptual challenge within the field of single-cell genomics^52^. We focus here on establishing methods for assessing consistent and accurate production of primary tissue cell-types at the class-level within organoids as a critical actionable first step towards increasing primary tissue fidelity across variable organoid differentiation protocols. While we prioritize broad cell-type comparisons, we also display the flexibility of our approach by scoring the preserved co-expression of finer resolution cell-type markers. This demonstrates our quantifications of preserved co-expression are applicable to a variety of cell-type annotation resolutions.

One exciting application for the use of neural organoid systems is the study of a wide-range of human neurological diseases using human *in vitro* models^53,54^, which critically depends on the *in vivo* fidelity of cell-types produced in organoids. Neural organoids have been used to model and investigate human disorders of neurodevelopmental^3,55^, neuropsychiatric^56–58^, and neurodegenerative^59–61^ nature, as well as infectious diseases^28,62,63^. It is essential that organoid systems model *in vivo* cell-types with extreme fidelity to fully realize the therapeutic potential of human organoids and ensure findings in these *in vitro* models are not specific to potential artifactual or inaccurate *in vitro* biology. While our results demonstrate that high primary tissue fidelity in organoids is currently methodologically possible, we also report a high degree of variability across organoids and studies/protocols indicating a remaining methodological gap. The broad applicability of our meta-analytic approach offers the potential for benchmarking primary tissue fidelity across numerous organoid protocols, aiding in increasing the quality of neural organoids for use in a wide-range of human health-related translational investigations.

The generalizable and flexible nature of our analysis is well suited to aid in the development of organoid differentiation protocols and the general quality control of neural organoids. Our results demonstrate the type of experiments possible through comparing preserved co-expression across organoid experimental variables, such as the differences in preserved co-expression between organoids grown in vertical or orbital shakers, as well as between transplanted or non-transplanted organoids. Importantly, our broad sampling across organoid protocols enabled clear identification of promising avenues for increasing organoid primary tissue fidelity. The strong performance across cell-types for the vascularized protocol we assessed suggests vascularized protocols as a route forward for global increases in primary fidelity. Additionally, our findings of specific ECM-related genes with consistent poorly preserved primary tissue co-expression in organoids suggests investigations into the interactions between Matrigel or other ECM-substrates and organoids may lead to general protocol adjustments for increasing primary tissue fidelity^64^. Looking beyond neural organoids, our framework for quantifying preserved co-expression can be applied to other organoid systems granted there is sufficient annotated primary tissue data to act as a reference.

## Methods

### Dataset download and scRNA-seq pre-processing

Links for all downloaded data (GEO accession numbers, data repositories, etc.) are provided in Supp. Table 1. All scRNA-seq data was processed using the Seurat v4.2.0 R package^65^. Data made available in 10XGenomics format (barcodes.tsv.gz, features.tsv.gz, matrix.mtx.gz) were converted into Seurat objects using the Read10X() and CreateSeuratObject() Seurat functions. Data made available as expression matrices were converted into sparse matrices and then converted into Seurat objects using the CreateSeuratObject() function. Ensembl gene IDs were converted into gene names using the biomaRt v2.52.0^66^ package.

Where metadata was made available, we separated data by batch (Age, Donor, Cell line, etc.) for our final total of 130 organoid and 51 primary tissue datasets (Supp. Table 1). We processed and analyzed each batch independently without integration. We used consistent thresholds for filtering cells across all datasets, keeping cells that had less than 50% of reads mapping to mitochondrial genes and had between 200 and 6000 detected genes. Several datasets provided annotations for potential doublets; we excluded all cells labeled as doublets when annotations were made available. All data made available with raw expression counts were CPM normalized with NormalizeData(normalization.method = ’RC’, scale.factor = 1e6), otherwise normalizations were kept as author supplied.

For primary tissue and organoid data made available with cell-type annotations, we provide our mapping between author provided annotations and our broad cell-type annotations in Supp. Table 2. Vascular annotated cell-types were excluded.

### Primary tissue MetaMarker generation and cross-validation

MetaMarkers were computed using the MetaMarkers v0.0.1^44^ R package, which requires shared cell-type and gene annotations across datasets to derive a ranked list of MetaMarkers. Gene markers for individual datasets were first computed using the compute_markers() function on the CPM normalized expression data for our annotated primary tissue datasets (Supp. Table. 1). A ranked list of MetaMarkers was then computed using the make_meta_markers() function using all 37 individual annotated primary tissue dataset marker lists. Genes are first ranked through their recurrent differential expression (the number of datasets that gene was called as DE using a threshold of log2 FC >= 4 and FDR p-value <= .05) and then through the averaged differential expression statistics of each gene across individual datasets. When we take the top 100 markers per individual dataset as in Fig. 2D, Fig. 5, Supp. Fig. 1A, and Supp. Fig. 3B, we rank markers for each dataset by their AUROC statistic as computed with the compute_markers() MetaMarkers function.

For the cross-validation of our primary tissue MetaMarkers, we excluded a single annotated primary tissue dataset, computed MetaMarkers from the remaining 36 annotated primary tissue datasets, and then used those MetaMarkers to predict the cell-type annotations of the left-out dataset. We construct an aggregate expression predictor to quantify the predictive strength a list of genes has, in this case our MetaMarker lists, in predicting cell-type annotations. Taking any arbitrary number of genes (10, 20, 50, 100, 250, or 500 MetaMarkers), we sum the expression counts for those genes within each cell and then rank all cells by this aggregate expression vector. We compute an AUROC using this ranking and the cell-type annotations for a particular cell-type through the Mann-Whitney U test. Formally:

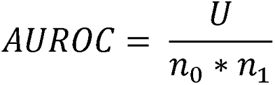

where U is the Mann-Whitney U test statistic, *n*_0_ is the number of positives (cells with a given cell-type annotation) and *n*_1_ is the number of negatives (cells without that cell-type annotation).

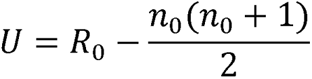

where *R*_0_ is the sum of the positive ranks.

As an example, if there are 10 genes that are perfect glutamatergic markers (only glutamatergic cells express these genes), then ranking cells by the summed expression of these genes will place all glutamatergic cells (positives) in front of all other cells (negatives), producing an AUROC of 1. The violin plots in Supp. Fig. 1B and in Figure 2E visualize our aggregate expression approach, where datapoints per cell-type are the aggregated expression counts for the given top 100 MetaMarkers across all cells per dataset (Supp. Fig. 1B) or aggregated across all datasets (Fig. 2E). We also compared the aggregate expression of the Neural Progenitor MetaMarkers across author provided cell-type annotations included in our broad Non-neuronal annotation, revealing the off-target expression of Neural Progenitor MetaMarkers is specific to annotated astrocytes (Supp. Fig. 1B).

For Supp. Fig. 1A, we took the top 100 cell-type markers per individual primary tissue dataset (x-axis) and used those genes to predict cell-type annotations as described above for all other annotated primary tissue datasets, reported as the AUROC boxplot distributions. The MetaMarker distribution was computed using a leave-one-out approach as described above. We ranked the individual primary tissue datasets by their median AUROC performance per cell-type to derive the distributions of ranks presented in Figure 2D, excluding the dividing progenitor data as performance was highly consistent across all primary tissue datasets.

### Cross-regional primary tissue MetaMarker expression

We investigated the aggregate expression of our top 100 MetaMarkers per cell-type across annotated brain regions separately for the annotated first-trimester and second-trimester primary tissue atlases due to differing regional annotations. MetaMarkers were computed with a leave-one-out approach as described above using all 37 of the annotated primary tissue datasets. For the heatmaps in Supp. Fig. 2, rows represent the annotated cells present within the given dataset, columns represent the aggregated expression for the top 100 given cell-type MetaMarkers and each annotated region present. We average the aggregated expression for each cell-type per region and then normalize each region (column) by the maximum average expression value across the cell-types. A value of 1 indicates that cell-type is the one maximally expressing the given MetaMarker set for that brain region. The heatmaps are ordered by cell-type and region and are not clustered.

### Organoid PCA

PCA analysis was performed using the Seurat function RunPCA() with the top 2000 variable features, determined using the Seurat function FindVariableFeatures(selection.method = ‘vst’, nfeatures = 2000). For each organoid dataset, we took the eigenvector for the first principal component, computed the absolute value, and then divided by the maximum value to compute a normalized vector between 0 and 1. We visualized the normalized eigenvectors for each organoid dataset in Supp. Fig. 3C, keeping primary tissue MetaMarker genes that were detected in the top 2000 variable genes of at least 10 organoid datasets. Genes missing from any given dataset’s top 2000 variable genes were given a value of 0. The heatmap was produced using the ComplexHeatmap v2.12.1^67^ package and was hierarchically clustered using the ward.D2 method for both rows and columns.

### Generating co-expression networks from scRNA-seq data

To generate a shared gene annotation space across all datasets, we fit each dataset to the GO gene universe before computing co-expression matrices. Using human GO annotations (sourced 2023-01-01 using the org.Hs.eg.db v3.15.0^68^ and AnnotationDbi v1.58.0^69^ R packages), we excluded gene expression from a dataset if the gene annotation was not present in GO and we zero-padded missing GO genes for each dataset.

We compute a gene-by-gene co-expression matrix per dataset using the spearman correlation coefficient computed across all cells in a given dataset. We then rank the correlation coefficients in the gene-by-gene matrix and divide by the maximum rank to obtain a rank-standardized co-expression matrix. All results reported using individual dataset co-expression networks (Fig. 3D-E, Fig. 4B, Figs. 5-6, Supp. Figs. 4-7) were obtained using the rank-standardized co-expression networks.

We compute the aggregated co-expression networks by taking the average of the rank standardized co-expression networks for each gene-gene index.

### Hierarchical clustering of primary tissue MetaMarkers by co-expression

We visualize the co-expression of primary tissue MetaMarker genes using the ComplexHeatmap package and the ward.D2 algorithm for hierarchical clustering. We use the fossil v0.4.0 package^70^ to compute the adjusted Rands Index with the adj.rand.index() function. To compute the adjusted Rands Index, we calculate a consensus clustering of MetaMarkers per co-expression network across 100 k-means clusterings (using the arguments row_km = 6, column_km = 6, row_km_repeats = 100, column_km_repeats = 100 within the Heatmap function) to compare to the perfect grouping of MetaMarkers by cell-type.

For the heatmaps in Fig. 3E, genes are ordered within each MetaMarker gene set by their average intra-gene set co-expression.

### Co-expression module learning analysis

EGAD v1.24.0^45^ is a machine learning framework that quantifies the strength of co-expression within an arbitrary gene-set compared to the rest of the genome with an AUROC quantification (Fig. 3C). We compute co-expression module AUROCs for all GO gene-sets (between 10 and 1000 genes per GO term) and our top 100 primary tissue MetaMarker gene-sets for each individual primary tissue and organoid co-expression network as well as the aggregated annotated, unannotated and organoid networks. For the annotated primary tissue co-expression networks, we employ a leave-one-out approach, learning MetaMarkers from 36 of the annotated datasets and computing co-expression module AUROCs for these MetaMarkers in the left-out dataset’s co-expression network. We compute co-expression module AUROCs using the EGAD run_GBA() function with default parameters. In Figure 3D, the ‘All GO terms’ distributions report the average co-expression module AUROC across all GO terms for each individual network.

### Preservation of co-expression

To compute our preservation of co-expression AUROC, we take the top 10 co-expressed partners for gene A in a reference co-expression network as our positive gene annotations. In a test co-expression network, we rank all genes through their co-expression with gene A and compute an AUROC using this ranking and the positive annotations derived from the reference network. If gene A in the test network has the exact same top 10 co-expressed partners as in the reference network, that would result in an AUROC of 1. To summarize a given gene-set’s preserved co-expression, we take the average preserved co-expression AUROC across all genes in that gene set as the preservation of co-expression score for that gene set. We use the aggregated annotated primary tissue co-expression matrix as our reference network.

The preserved co-expression scores for the annotated primary tissue data in Figure 4B were computed with a leave-one-out approach. MetaMarkers and an aggregated co-expression matrix were computed from 36 of the annotated primary tissue datasets and then preserved co-expression scores were computed using the co-expression network of the left-out annotated primary tissue dataset.

### Preservation of fine resolution cell-types

To define markers for finer resolution cell-types, we utilize the differential expression (DE) statistics computed from a study that performed a morphogen screen in neural organoids and reported extensive neural cell-type diversity^48^. For each cell-type, we rank genes by their adjusted DE p-value and take the top 10 genes per cell-type to compute preserved co-expression scores. When comparing against our MetaMarker gene sets in Figure 4E, we ensure no overlap in the top 10 cell-type and top 100 MetaMarker gene sets.

### Preservation of GO term co-expression

We compute p-values for the preservation of co-expression of GO terms using a mean sample error approach. Using the aggregated annotated primary tissue co-expression network as the reference and the aggregated organoid network as the test network, we first compute the preserved co-expression AUROCs for all individual genes, taking the mean and standard deviation value as the population mean and population standard deviation. For any given GO term, we first compute the preserved co-expression score for the term (the average of the preserved co-expression AUROCs for the genes in the term) and then compute the sample error for that score with:

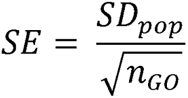

where *SD*_*pop*_is the population standard deviation and *n*_*GO*_ is the number of genes in the GO term. We then compute a z-score through:

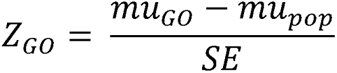

where *mu*_*go*_ is the preserved co-expression score for the GO term and *mu*_*pop*_ is the population mean preserved co-expression AUROC. We compute left-sided p-values using the standard normal distribution:

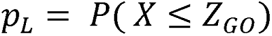

Where X is a normal distribution with mean = 0 and standard deviation = 1. We use the R function pnorm(*z*_*GO*_) to compute this p-value.

We then compute the right-sided p-value as:

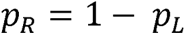

We adjust p-values using the R function p.adjust(method = ‘BH’). We filter for GO terms that have between 20 and 250 genes per term and use a threshold of FDR-corrected p-value <= 0.0001 to call significance. Significant left-sided p-values are interpreted as GO terms with significantly smaller preserved co-expression scores (significantly not preserved) than expected through sampling error and right-sided p-values are interpreted as GO terms with significantly larger preserved co-expression scores (significantly preserved) than expected through sampling error. We use the R package rrvgo to visualize the significant GO terms in Fig. 4F.

### Computing correlation significance

We employ a permutation test to compute p-values for any given correlation coefficient. We permute data-pairs and compute a correlation coefficient, repeating for 10,000 random permutations to generate a distribution of correlation coefficients under the null hypothesis of independence. We calculate a two-sided p-value for the original correlation coefficient as the number of permuted correlation coefficients whose absolute value is greater than or equal to the absolute value of the original correlation coefficient, divided by 10,000. We adjust p-values using the R function p.adjust(method = ‘BH’) and use a FDR-corrected p-value threshold of <= .05 to call significance.

### Comparing co-expression of normal vs. perturbed organoids

For both the co-expression module AUROCs and the preserved co-expression scores of normal and perturbed organoids, we test for significant differences per cell-type using the Mann Whitney U test, adjusting p-values with the R function p.adjust(method = ‘BH’) and using a FDR-corrected p-value threshold of <= .05 to call significance.

### Organoid temporal analysis

The organoid temporal analysis for both predicting organoid annotations with primary tissue markers (Supp. Fig. 3B) and scoring the preserved co-expression of organoid co-expression using primary tissue networks as reference (Fig. 5) were performed for all pair-wise combinations of the 37 annotated primary tissue datasets and the 26 temporally annotated forebrain organoid datasets. We excluded the GW7-28 annotated primary tissue dataset from the temporal preserved co-expression analysis (Fig. 5) due to the wide temporal range sampled. For predicting organoid annotations with primary tissue markers, we used the top 100 markers per primary tissue dataset to construct aggregate expression predictors in the organoid datasets as described above. The MetaMarkers performance was calculated using MetaMarkers derived from all 37 annotated primary tissue datasets. For scoring preserved co-expression, individual primary tissue networks were used as the reference with individual organoid networks as the test networks. We computed the preserved co-expression scores of the top 100 primary tissue cell-type markers per individual primary dataset for each individual organoid network.

### GO enrichment analysis

We compute enrichment for GO terms using Fisher’s Exact Test as implemented through the hypergeometric test. We compute raw p-values for GO terms with between 10-1000 genes and compute FDR-adjusted p-values using p.adjust(method = ‘BH’). We only consider GO sets with between 20 and 500 when choosing the top 10 GO sets in Figure 4G, ranked by FDR-adjusted p-value.

### R and R packages

All analysis was carried out in R v4.3.1. Colors with selected using the MetBrewer v0.2.0 R library. Plots were generated using ggplot2 v3.3.6^71^. Spearman correlation matrices for co-expression networks were computed using a python v3.6.8 script, implemented in R with the reticulate v1.26 R package, as well as using functions from the matrixStats v0.62.0 R library. All code used in generating results and visualizations will be made public at the time of publication. The preservedCoexp R library is made available at https://github.com/JonathanMWerner/preservedCoexp. All code used for analysis is made available at https://github.com/JonathanMWerner/meta_organoid_analysis.

## Author Contributions

JMW and JG conceived the project. JMW and JG designed analyses. JMW performed analyses. JMW and JG wrote the manuscript.

## Acknowledgments

JMW was supported by NSF award no. DGE-1938105. JG and JMW were supported by NIH grants R01MH113005 and R01LM012736. This material is based upon work supported by the National Science Foundation Graduate Research Fellowship Program under grant no. DGE-1938105. Any opinions, findings, and conclusions or recommendations expressed in this material are those of the author(s) and do not necessarily reflect the views of the National Science Foundation. We thank Tomasz Nowakowski and members of the Gillis lab for helpful comments on the manuscript.

