## Supplemental Figures for "Preservation of co-expression defines the primary tissue fidelity of human neural organoids"

**Supplemental data**

**Supplemental Table 1**

Table containing the study origin and download links for all primary tissue and organoid scRNA-seq datasets. The batch variable column details the meta-data used in determining batch. The region/protocol column details the sampled primary tissue brain regions or the organoid differentiation protocol.

**Supplemental Table 2**

Table containing our mapping between author provided annotations (Author annotations column) and our broad cell-type annotations (Class annotations column).

**Supplemental Figure 1**

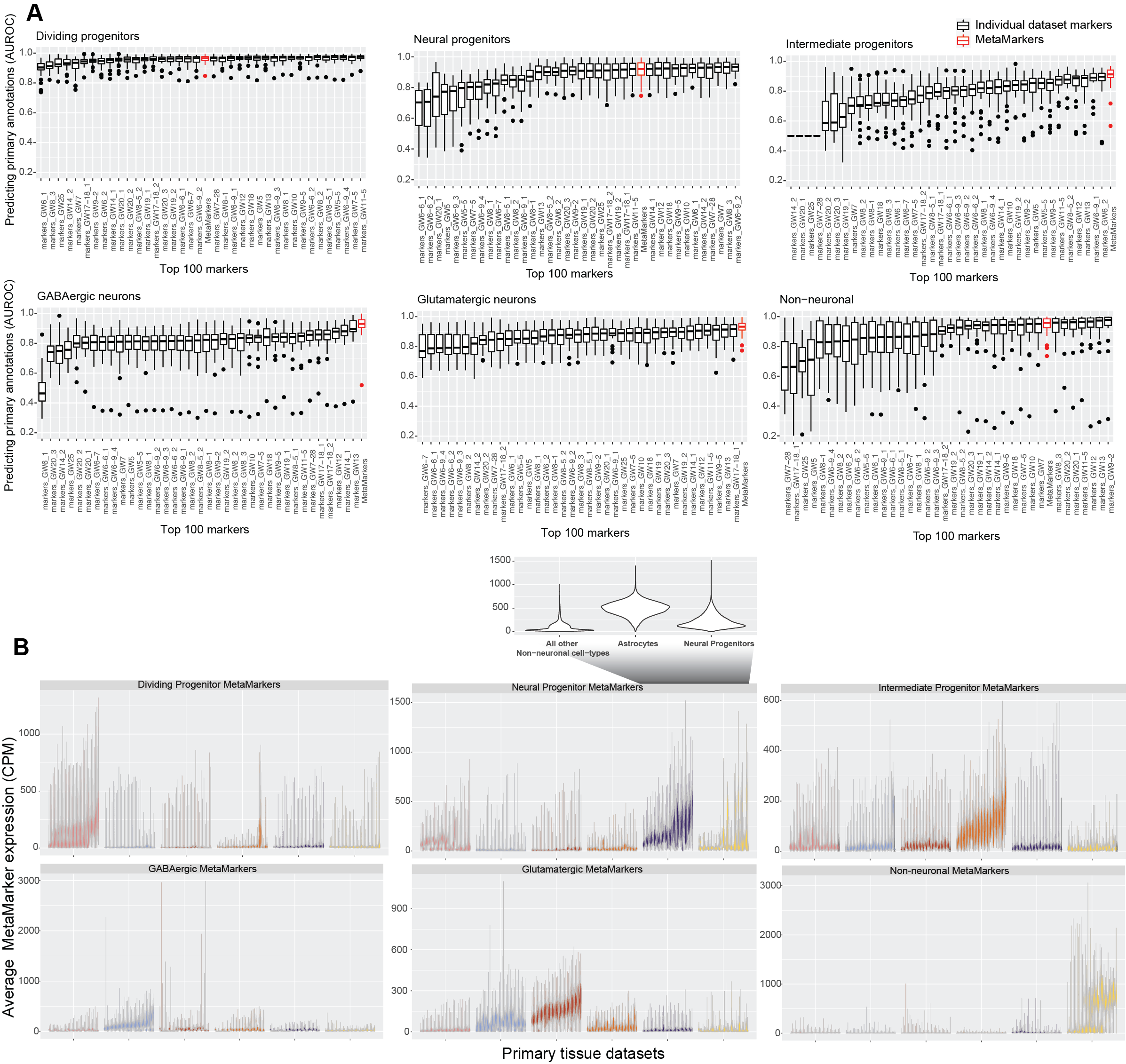

**MetaMarkers as temporally robust primary tissue cell-type markers**

**A** MetaMarkers are consistent top performers in predicting primary tissue cell-type annotations. Boxplots of AUROCs for predicting cell-type annotations across all primary tissue datasets using the top 100 marker genes per individual primary tissue dataset compared to MetaMarkers (red). Datasets are ordered by their median performance, providing the rank distributions in Figure 2D.

**B** MetaMarkers exhibit cell-type specificity across all primary tissue datasets. Averaged distributions of gene expression for the top 100 MetaMarkers across all annotated primary tissue datasets with leave-one-out cross-validation. Figure 2E is the aggregate over these individual dataset distributions. Inset displays the average Neural Progenitor MetaMarker expression for Neural Progenitor, Astrocyte, and all non-astrocyte Non-neuronal cells

**Supplemental Figure 2**

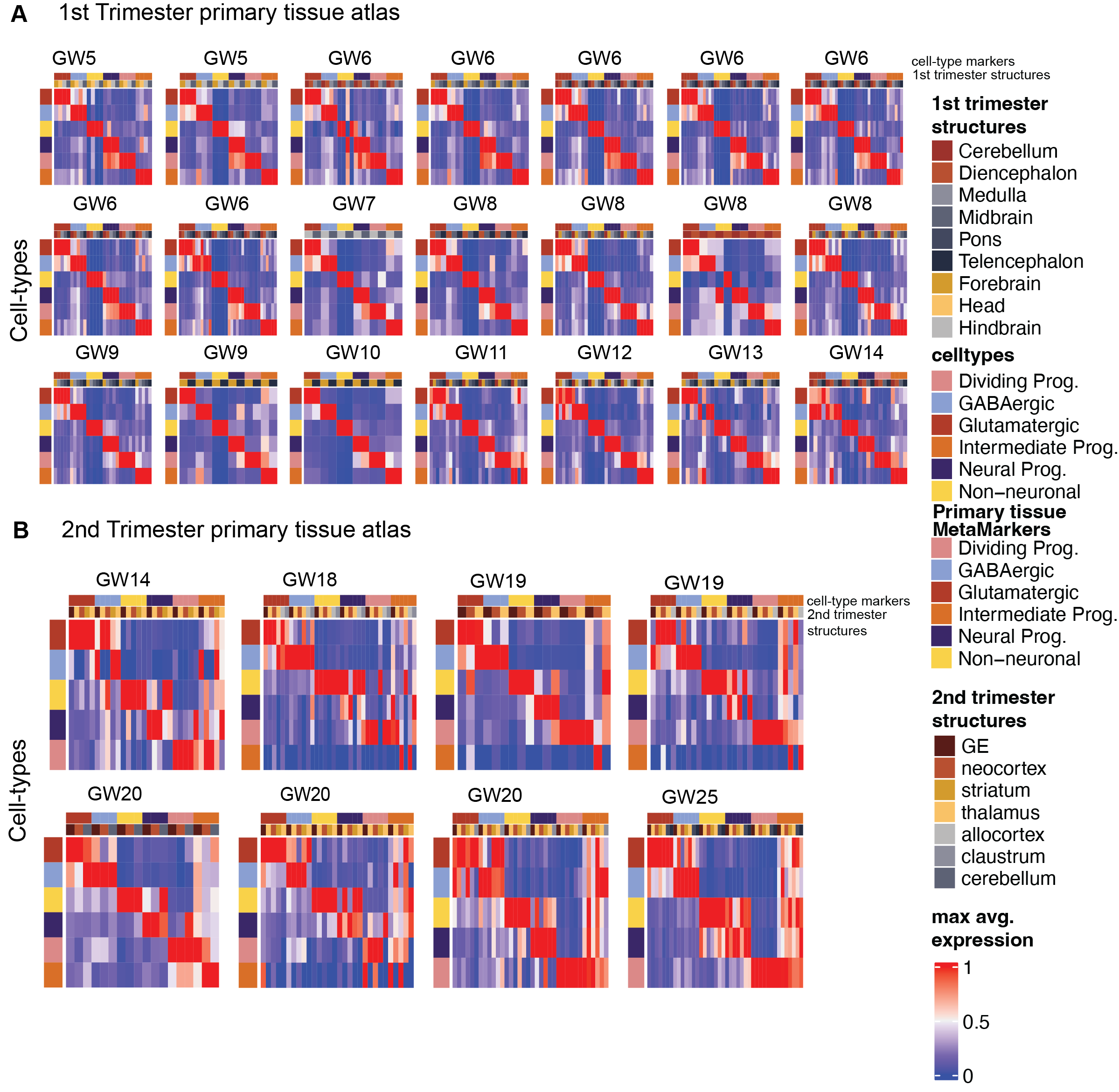

**MetaMarkers as regionally robust primary tissue cell-type markers**

**A** MetaMarkers exhibit cross-regional cell-type specificity. Heatmaps of maximum normalized average MetaMarker expression for cell-types and brain regions of the first trimester annotated primary tissue atlas. Cell-types comprise the rows with MetaMarker gene expression for cells from each annotated brain region comprising the columns. Data is maximum normalized per region/column.

**B** MetaMarkers exhibit cross-regional cell-type specificity. Heatmaps of maximum normalized average MetaMarker expression for cell-types and brain regions of the second trimester annotated primary tissue atlas. Cell-types comprise the rows with MetaMarker gene expression for cells from each annotated brain region comprising the columns. Data is maximum normalized per region/column.

**Supplemental Figure 3**

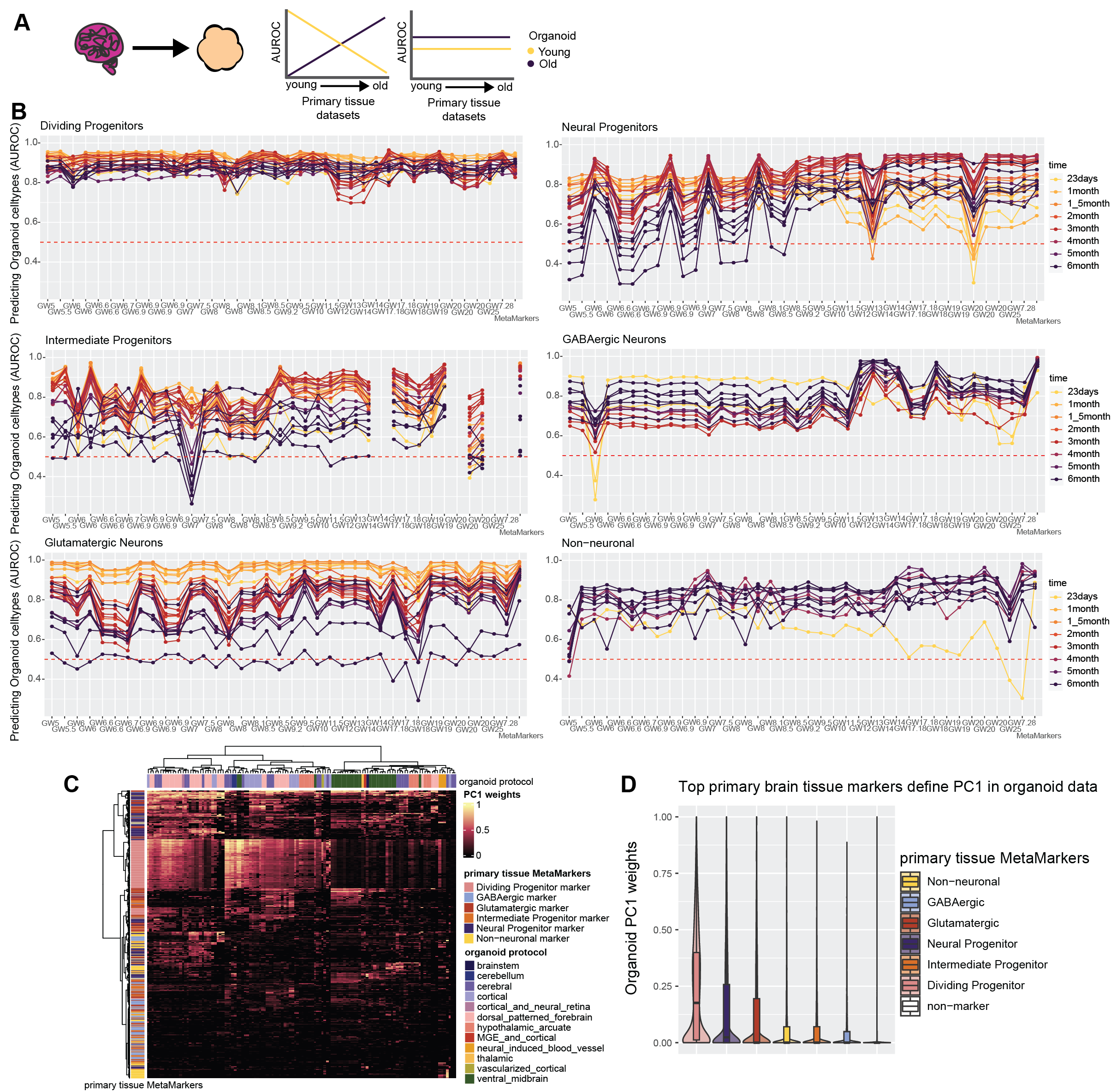

**Primary tissue MetaMarkers consistently predict organoid cell-types across timepoints**

**A** Schematic showing two potential outcomes when comparing cell-type marker expression between primary tissue and organoid data on a temporal axis. There may be a temporal relationship, with younger organoids recapitulating younger primary tissue marker expression over older primary tissue marker expression and vice versa for older organoids, or there may be no temporal relationship.

**B** Broad primary tissue cell-type markers have consistent performance predicting organoid annotations independent of temporal variation. Line plots showing the cell-type prediction AUROCs using top 100 markers from individual primary tissue datasets for all organoid time points. Primary tissue datasets on the x-axis are ordered from youngest to oldest.

**C** Primary tissue MetaMarkers define the first organoid principal component. Heatmap of normalized eigenvalues for primary tissue MetaMarkers within the first principal component of each organoid dataset.

**D** MetaMarker gene-set distributions of normalized PC1 eigenvalues across all organoid datasets.

**Supplemental Figure 4**

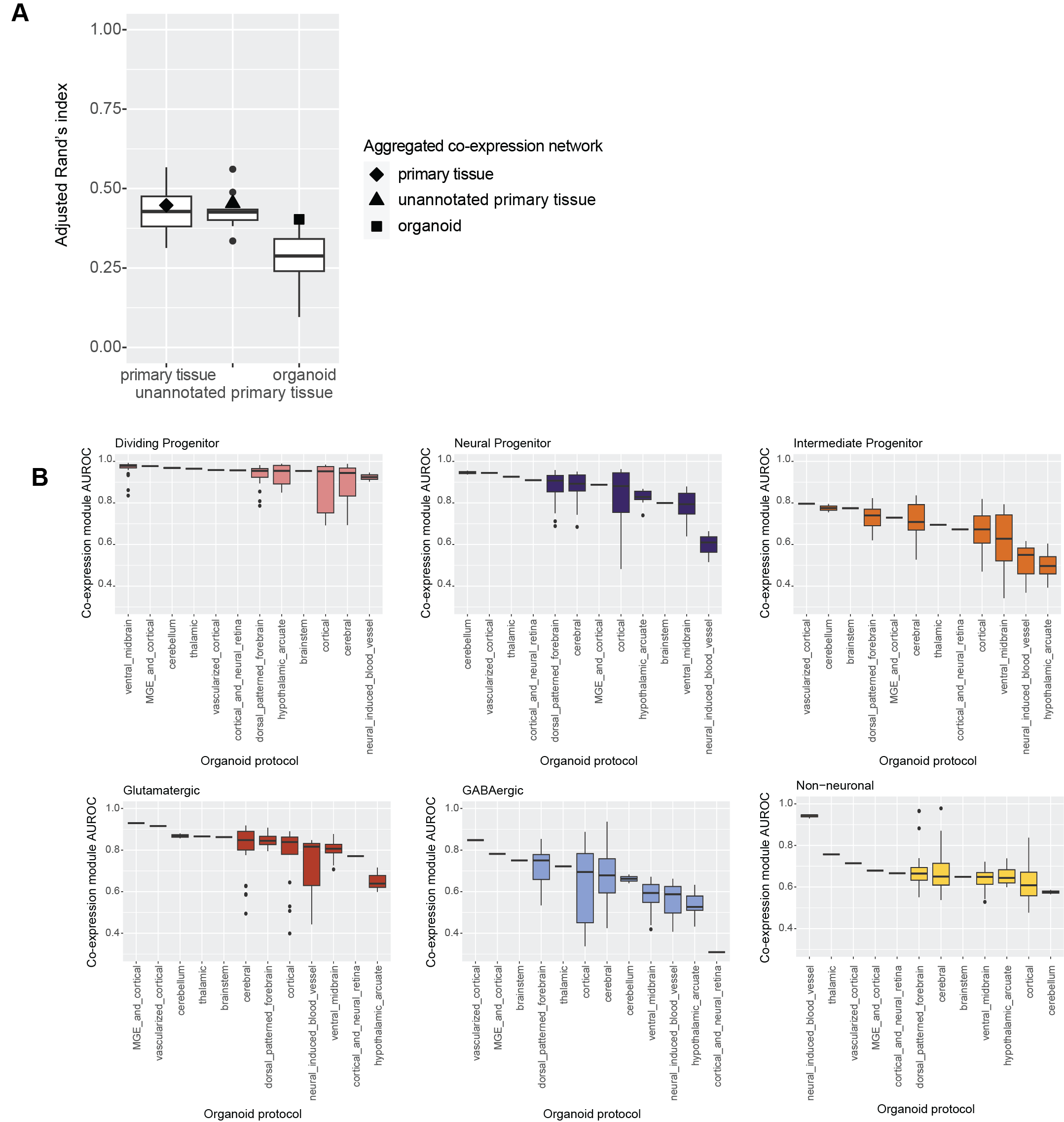

**Intra-marker set MetaMarker co-expression varies over organoid protocols**

**A** Organoid cell-type clustering via co-expression is notable lower compared to all primary tissue datasets. Distributions of the Adjusted Rands Index (ARI) for individual annotated primary tissue, unannotated primary tissue, and organoid datasets. The ARI scores for the aggregate networks are denoted with the special characters.

**B** Organoids vary by protocol type for their primary tissue cell-type co-expression module scores. Boxplot distributions of co-expression module scores for the primary tissue MetaMarkers computed from organoid co-expression networks. Scores for organoid networks are grouped by organoid protocol type and ordered by their median score.

**Supplemental Figure 5**

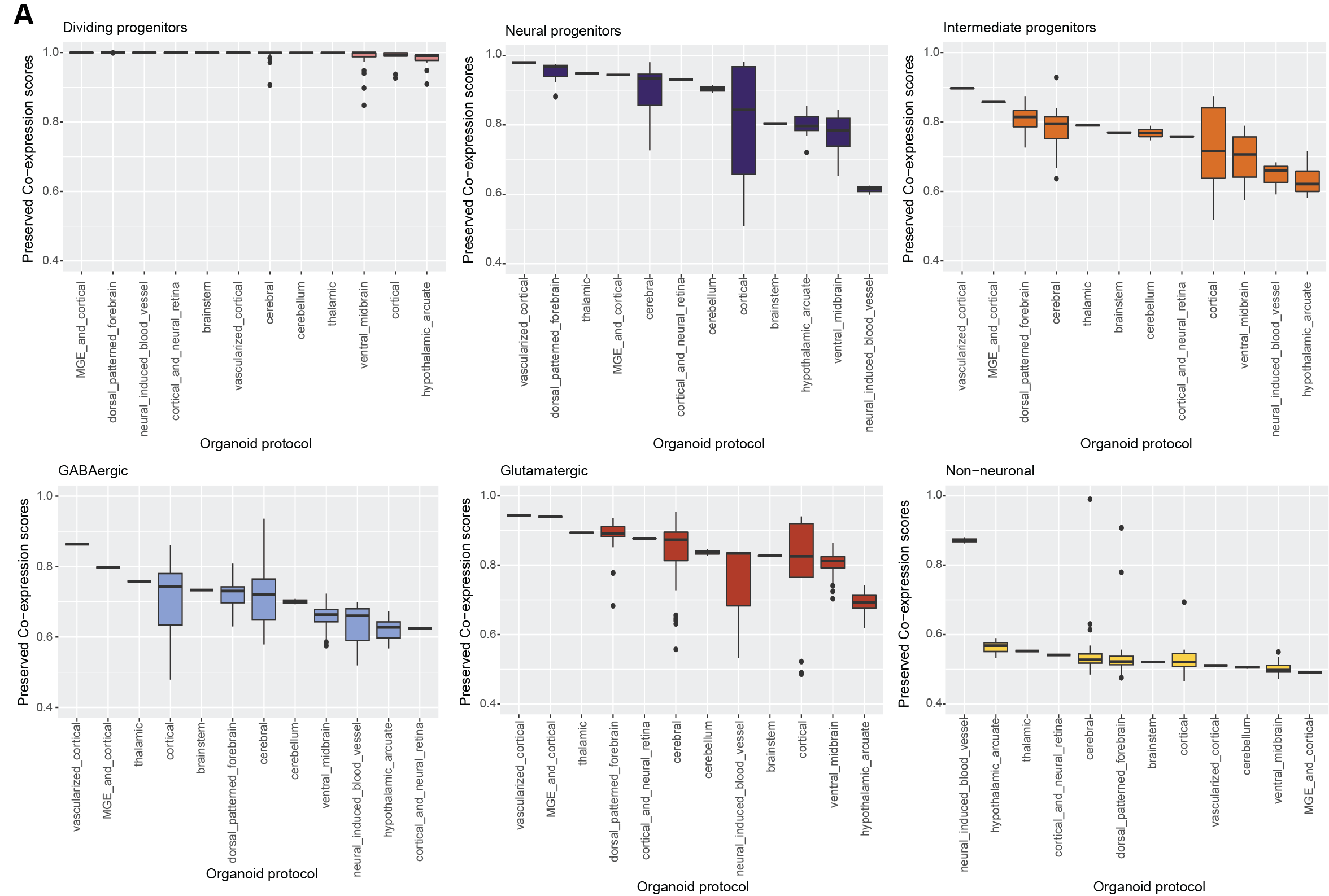

**Preservation of MetaMarker set co-expression varies over organoid protocols**

**A** Organoids vary by protocol type for their primary tissue cell-type preserved co-expression scores. Boxplot distributions of preserved co-expression scores for the primary tissue MetaMarkers computed from organoid co-expression networks. Scores for organoid networks are grouped by organoid protocol type and ordered by their median score.

**Supplemental Figure 6**

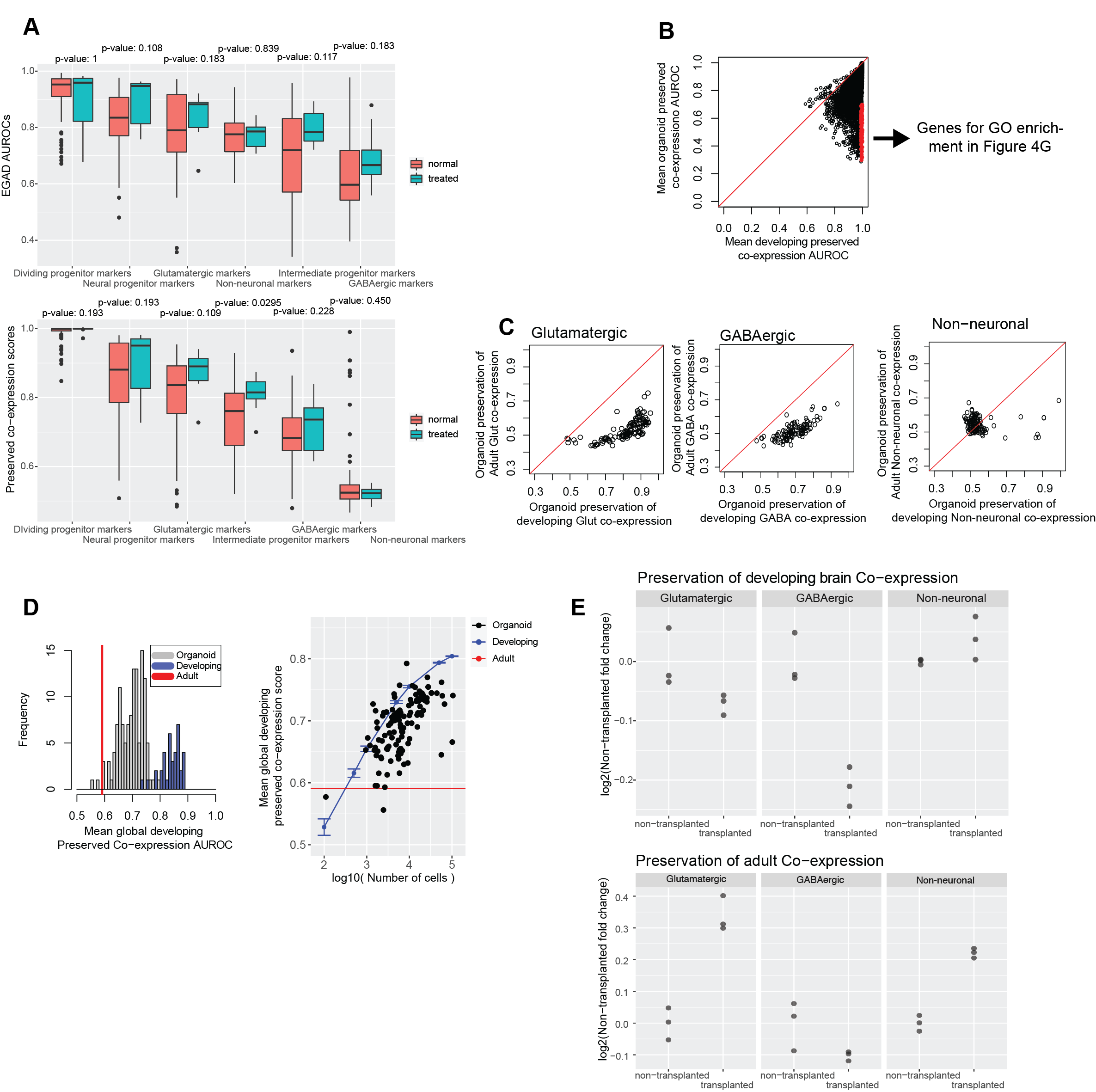

**Neural organoids preserve co-expression of developing neural tissue over adult neural tissue**

**A** Normal and treated organoids exhibit no differences in their recapitulation of primary tissue co-expression. Boxplots comparing either the co-expression module scores or preserved co-expression scores by cell-type between normal and treated organoids.

**B** Organoids globally have low preserved developing brain co-expression of individual genes across the genome. Points show the average preserved developing brain co-expression AUROC of individual genes, comparing the average across developing brain networks (x-axis) against the average across organoid networks (y-axis). The points colored in red are genes with developing brain scores >= 0.99 and organoid scores < 0.70.

**C** Organoids preserve developing neuronal co-expression over adult co-expression. Scatterplots showing the preserved co-expression scores of either the top 100 developing brain MetaMarkers (x-axis) or the top 100 adult MetaMarkers (y-axis).

**D** Organoids lie between adult and developing brain data for global preservation of developing brain co-expression. Distributions of average preserved developing brain co-expression AUROCs across all genes for organoid and developing brain networks. The redline shows the performance of the adult co-expression network. The scatterplot plots the data in the histogram (y-axis) against the number of cells in each organoid dataset (x-axis). The blue line shows performance for a cell down-sampled developing brain dataset, with points representing the average performance over 10 random samples and the error bars showing ± 1 standard deviation.

**E** Transplanted organoids preserve adult co-expression over developing brain co-expression. Points represent the log2-fold change over the mean performance of the non-transplanted organoids for preserved co-expression scores.

**Supplemental Figure 7**

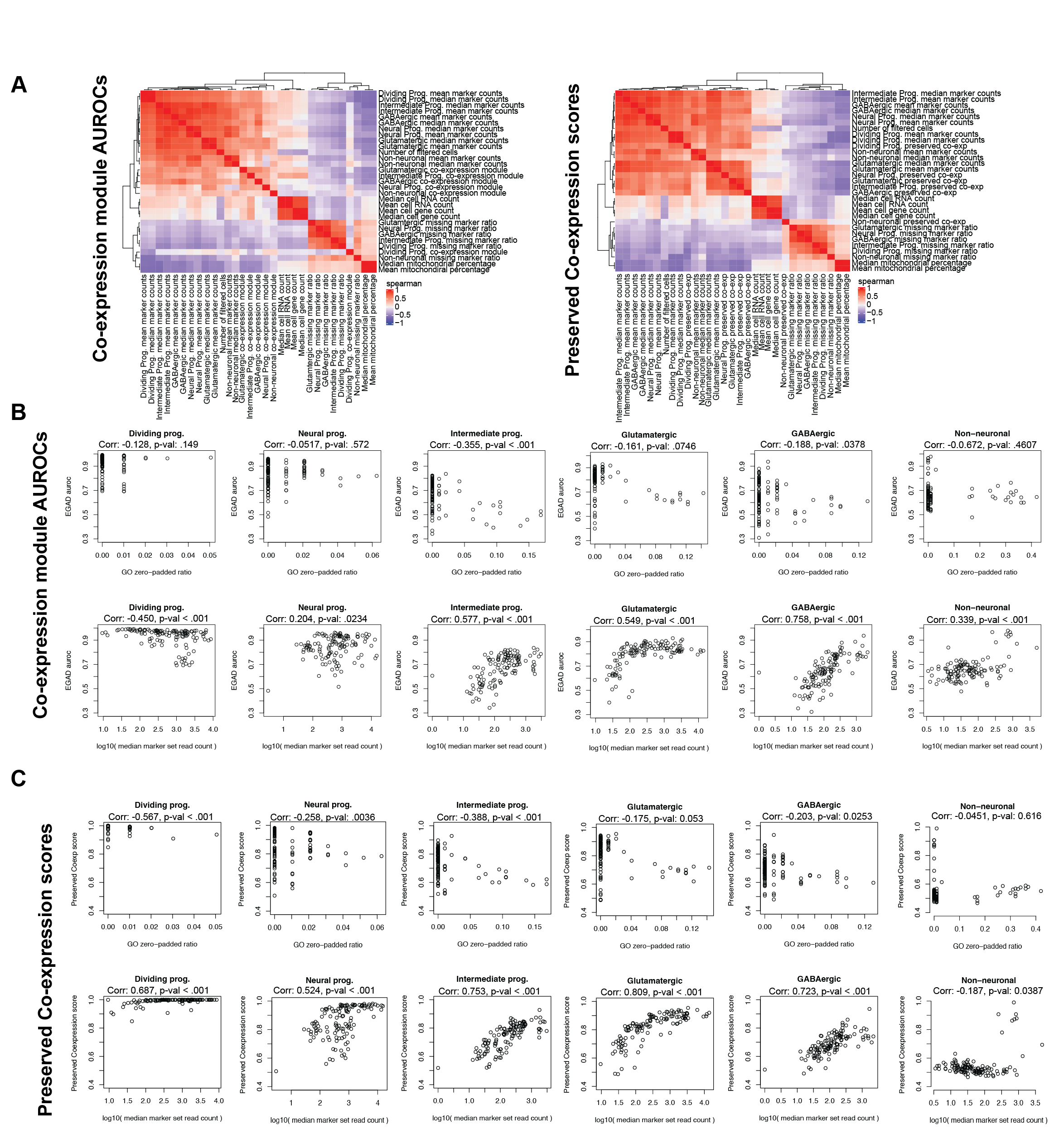

**Strength of MetaMarker co-expression in organoids is related to expression levels**

**A** Marker set expression and cell number are strongly correlated with co-expression performance across organoid datasets. Heatmaps of spearman correlations between either co-expression module scores or preserved co-expression scores and various technical features of each network/dataset, like marker set expression, dataset sequencing depth, number of cells in each dataset, and the zero-padding ratio of each marker set.

**B** Scatterplots of either the zero-padded ratio (top row) or marker set expression (bottom row) against the co-expression module scores for each cell-type across the organoid datasets.

**C** Scatterplots of either the zero-padded ratio (top row) or marker set expression (bottom row) against the preserved co-expression scores for each cell-type across the organoid datasets.

**References** (Supp. Table 1)
